# Saturation mutagenesis genome engineering of infective ΦX174 bacteriophage via unamplified oligo pools and golden gate assembly

**DOI:** 10.1101/798546

**Authors:** Matthew S. Faber, James T. Van Leuven, Martina M. Ederer, Yesol Sapozhnikov, Zoë L. Wilson, Holly A. Wichman, Timothy A. Whitehead, Craig R. Miller

## Abstract

Here we present a novel protocol for the construction of saturation single-site—and massive multi-site—mutant libraries of a bacteriophage. We segmented the ΦX174 genome into 14 non-toxic and non-replicative fragments compatible with golden gate assembly. We next used nicking mutagenesis with oligonucleotides prepared from unamplified oligo pools with individual segments as templates to prepare near-comprehensive single-site mutagenesis libraries of genes encoding the F capsid protein (421 amino acids scanned) and G spike protein (172 amino acids scanned). Libraries possessed greater than 99% of all 11,860 programmed mutations. Golden Gate cloning was then used to assemble the complete ΦX174 mutant genome and generate libraries of infective viruses. This protocol will enable reverse genetics experiments for studying viral evolution and, with some modifications, can be applied for engineering of therapeutically relevant bacteriophages with larger genomes.

## Introduction

Predicting the tempo and trajectory of evolutionary change in the complex environments encountered by viruses and bacteria remains a challenge (Holmes 2013, Neher 2013, Lieberman et al. 2014). Understanding such changes are important for fundamental evolutionary studies as well as biotechnology applications like phage therapy for multi-drug resistant bacteria (Dedrick et al. 2019). Two obstacles preventing better predictions are the oversimplification of the environment in experimental evolution studies compared to wild conditions and the vast number of experimentally unexplored mutational combinations that can occur, even in short adaptive walks and small genomes. Advances in DNA sequencing and synthesis have opened new ways to study microbial evolution and overcome some of these obstacles (Barrick et al. 2013, Fowler et al. 2014). Unfortunately, suitable methods do not exist for generating comprehensive bacteriophage mutant libraries. This technology gap hinders those seeking to engineer phage for biotechnological applications and for those seeking a deeper understanding of how viruses evolve.

Methods for genetically engineering phages have recently been reviewed (Pires et al. 2016) and commonly involve homologous recombination and recombineering. While such approaches can be improved by clever incorporation of CRISPR-Cas systems (Kiro et al. 2014, Lemay et al. 2017, Schilling et al. 2018), the overall modest efficiencies largely limits mutagenesis to the generation and single-site mutation of chimeric genomes (Doore and Fane 2015, Doore et al. 2017), gene deletions (Uchiyama et al. 2009), and limited multi-site mutagenesis (Stockdale et al. 2015). A recent approach combines mutagenesis by PCR with degenerate (NNK) primers with in-cell recombination to create variation at specific residues in the tail fiber of T3 phage (Yehl et al. 2019). By contrast, human viruses often have reverse genetics systems in place where comprehensive mutant libraries can be prepared for single genes (Lee et al. 2018) or regions within a gene (Doud et al. 2018, Wu and Qi 2019) using replicative plasmids that encode whole viruses or viral components. Similarly, deep mutational scanning can be performed on plasmid-encoded phage proteins like the MS2 capsid protein (Hartman et al. 2018). As described by Hartman et al. 2018, such experiments can identify mutations that confer useful characteristics, such as MS2 capsid stability mutations that could improve the targeted delivery of therapeutics to particular cellular compartments.

Advances in the technologies for generating mutant libraries, synthesizing DNA, and for the assembly of large DNA fragments (van Dolleweerd et al. 2018) allow for the construction and assembly of user-defined mutagenesis of long nucleic acids. Nicking mutagenesis (NM) can be used to construct comprehensive single-site or other user-defined mutant libraries (Wrenbeck et al. 2016) using dsDNA as a template. In NM, oligos encode the desired mutations by mismatch with the parental template. On-chip ink-jet printed oligo pools have been integrated into NM (Medina-Cucurella et al. 2019), so one can now construct heterogeneous libraries of oligos that contain tens to hundreds of thousands of high fidelity, unique sequences (Kosuri and Church 2014, Medina-Cucurella et al. 2019) with a low per base pair cost.

We therefore suspected that large user-defined libraries of phage could be generated by combining the advantages of replicative plasmids with advances in DNA synthesis. The bacteriophage ΦX174 is an excellent system to test this approach. High resolution X-ray crystallography structures of the capsid and spike proteins (McKenna et al. 1992, Dokland et al. 1997) are available for ΦX174, making it amenable to compare empirical mutagenesis results and molecular modeling predictions, it has a very small genome (Sanger et al. 1978) of 5,386 nucleotides that is readily sequenced, and it is easily grown on a variety of lab *E. coli* strains. These attractive features of the ΦX174 experimental system have led to proof-of-principle demonstrations from other groups for synthetic genome assembly from oligonucleotides (Smith et al. 2003) and complete refactoring of the phage (Jaschke et al. 2012, Jaschke et al. 2018).

ΦX174 has also been extensively studied (Aoyama et al. 1981, Hafenstein and Fane 2002, Bernal et al. 2004, Rokyta et al. 2006), especially as a model system for experimental evolution (Bull et al. 1997, Crill, Wichman, and Bull 2000, Wichman, Millstein, and Bull 2005) since populations grow rapidly to very large size, mutation rate is high, and the fitness differences between even single mutations can be very large. One of the focal areas of work on ΦX174 and related microvirid phage has been testing models of adaptive evolution and epistasis with empirical data (Rokyta et al. 2005, Caudle, Miller, and Rokyta 2014, Sackman and Rokyta 2018). The inferential power of these studies has been limited by the modest sizes of the mutational libraries.

Here we have combined Golden Gate assembly, nicking mutagenesis, and oligo pool technology to construct near comprehensive single-site and expansive multisite mutant libraries for the genes encoding the capsid F protein and spike G protein of the bacteriophage ΦX174. To our knowledge this is the first time near comprehensive single site mutant libraries of full capsid and spike proteins have been generated for an infective bacteriophage.

## Results

Our strategy for generating large user-defined mutagenesis libraries of bacteriophage ΦX174 is shown in Figure 1. The circular 5386 nucleotide genome compactly encodes 11 genes where the F and G genes encode the capsid and spike proteins, respectively (Fig. 1A). The genome was segmented onto 14 separate plasmids, including 3 plasmids for the F gene and 2 for the G gene (Fig. 1B). Fragment boundaries were first engineered to divide large genes, separate regulatory regions from protein coding genes, and result in fragments of similar size. Initial attempts to clone some fragments resulted in low plasmid yield, presumably due to gene toxicity. These genes were further divided and tested empirically, resulting in the final set of 14 fragments. User-defined comprehensive mutations on F and G were encoded using nicking mutagenesis with a single unamplified oligo pool containing all 11,860 mutagenic oligonucleotides (Fig. 1C). ΦX174 mutant genomes were reconstituted from individual plasmids using Golden Gate cloning and transformation into a host *E. coli* strain (Fig. 1D). Harvested phages contained all possible non-synonymous mutations in the F and G genes depending on the mutagenized segments used for Golden Gate cloning (Fig. 1E).

**Figure 1.**
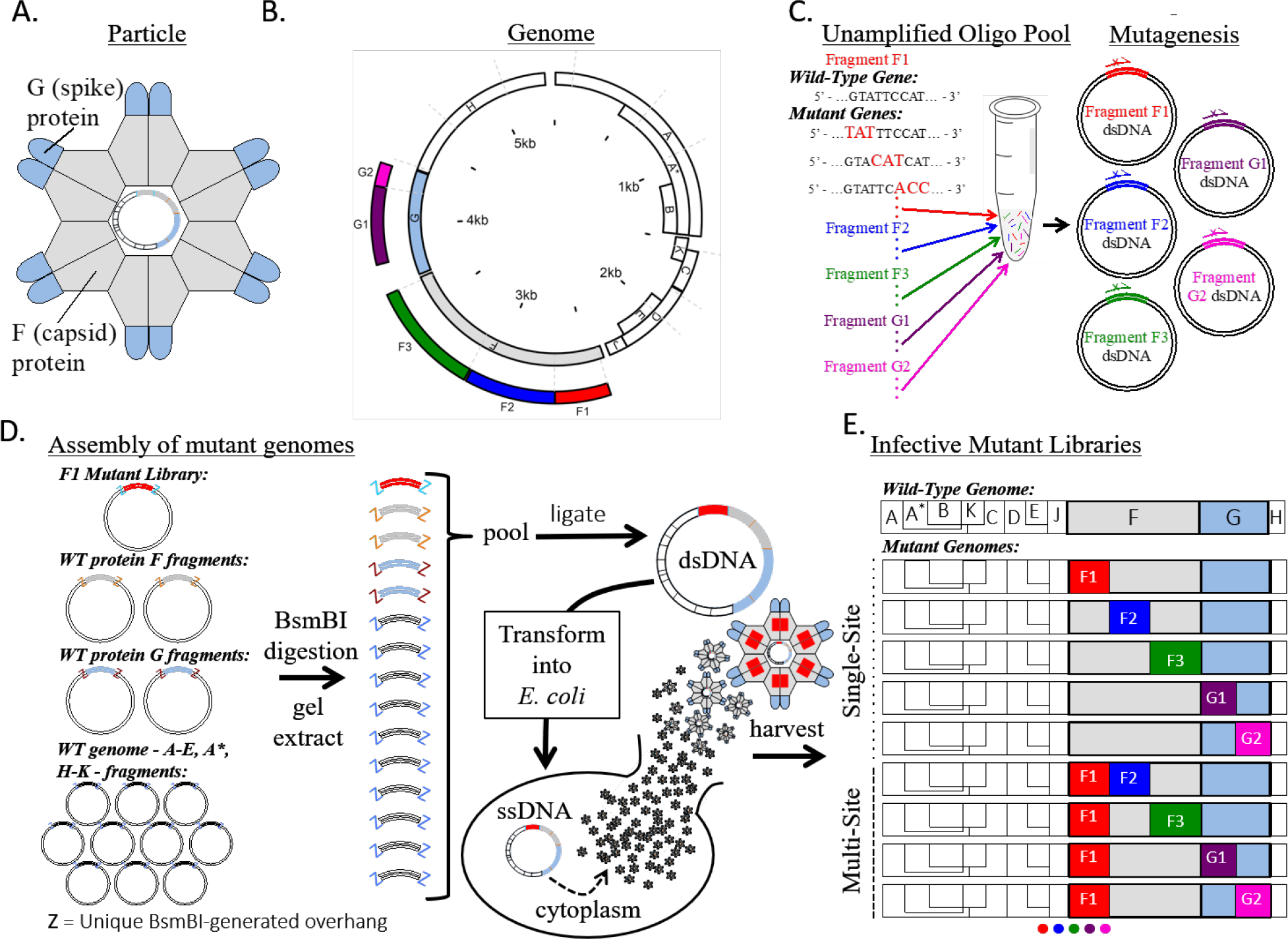
ΦX174 mutant library assembly. **A.** Cartoon structure of the ΦX174 virion. F and G proteins are organized into pentamers and are the only exposed proteins in the mature virion. **B.** A linear schematic of the circular ΦX174 genome showing the size and location of the synthetic genome fragments. The location of mutated residues in genes F and G are indicated. **C.** An oligo pool containing all mutagenic oligos for genes F and G was generated and used in Nicking Scanning Mutagenesis for the generation of saturation mutant libraries for genes F and G. **D.** Golden Gate cloning was used to assemble the mutant genomes, which were transformed into XL1-Blue competent cells (Agilent) and plated on susceptible *E. coli* C cells. The resulting plaques were sequenced. **E.** Linear schematics depicting some of the possible mutant libraries that can be generated using the workflow and existing libraries.

We sought a reverse genetics system for ΦX174 wherein the virus chromosome was segmented and encoded on individual vector plasmids. Our method of construction closely followed that for the assembly of human coronavirus NL63 (Donaldson et al. 2008), where each plasmid contained unique BsmB1 type IIS restriction endonuclease sites flanking unique five nucleotide overlaps of wild-type (WT) ΦX174. This architecture allows for faithful and easy assembly of the complete genome via Golden Gate cloning. The ΦX174 genome was separated into 14 separate nontoxic fragments by PCR and cloned into vector plasmids where genes were separated from their promoters and larger genes were segmented (full sequences for all plasmid inserts are given in Supplemental Note S1). The segments ranged from 129 to 634 base pairs in length, with necessary additional truncations of genes D and A. ΦX174 phage could be reconstituted by digesting pooled plasmids with BsmB1, ligating inserts overnight, and transforming into electrocompetent *E. coli* C cells.

The F and G genes were targeted for saturation mutagenesis. We introduced mutations by nicking mutagenesis, which requires the presence of a BbvCI nicking site on the dsDNA plasmid. The F gene plasmid F3 encoding residues 246-427 of the F gene product contained a unique BbvCI sequence, while the remaining four plasmids (F1, F2, G1, G2) required introduction of BbvCI nicking sites in the vector backbone. The presence in each plasmid of the unique BbvCI nicking site was verified with the successful generation of circular ssDNA from the modified plasmids by BbvCI.Nt and exonuclease digestion (Supplemental Figure S1).

In nicking mutagenesis, desired mutations are introduced by libraries of mutagenic oligonucleotides, which can be sourced from unamplified oligo pools (Medina-Cucurella et al. 2019). We synthesized a custom pool of 11,860 oligos that encoded nearly every missense and nonsense mutation in the F and G genes. Each missense or nonsense mutation was encoded by a single oligonucleotide. Application of nicking mutagenesis using this oligo pool for different plasmids (F1, F2, F3, G1, G2) resulted in at least 14-fold excess transformants required for 99.9% theoretical coverage of the desired library (Supplemental Table S3). The diversity of the mutant plasmid libraries was validated using deep sequencing on an Illumina MiSeq platform. All plasmid libraries have >99.7% coverage of all possible single mutations (n = 11,821/11,860, minimum counts for a mutation >5, full library statistics are given in Table 1 and Supplemental Table S4). Heatmaps showing mutation-specific frequencies for all segments are shown in Supplementary Figures S2–S9.

**Table 1.**
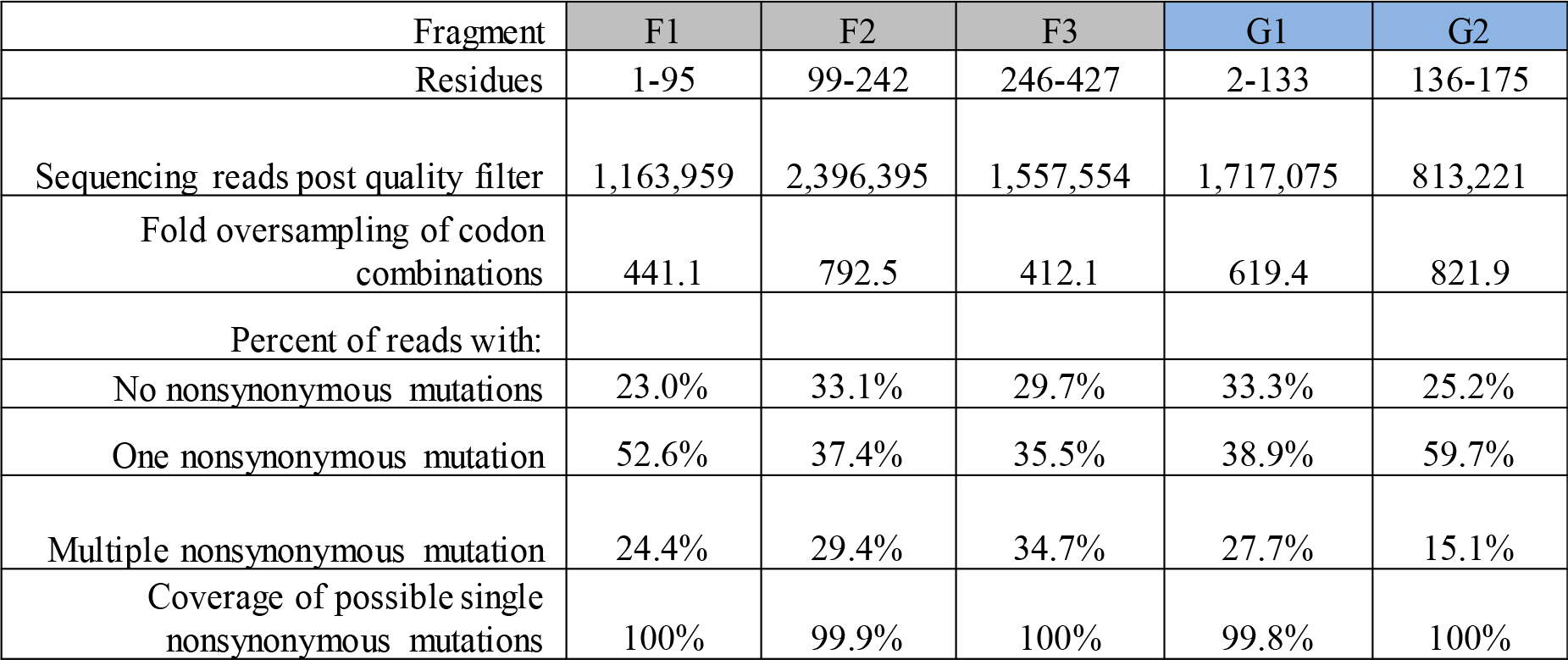
Mutant library NGS statistics. Summary table of the of the plasmid libraries prior to viral genome assembly.

Mutant dsDNA libraries were transformed into non-susceptible hosts (Agilent XL1-Blue), then plated on susceptible hosts (*E. coli* C) after a 30-minute recovery in SOC. Viral titers increase rapidly after 30 minutes, suggesting that the XL1-Blue cells do not burst until after this time point (Supplemental Table S5). Accordingly, we reasoned that the number of plaques at 30 minutes represents the number of XL1-Blue cells that were transformed with viable genomes. The number of transformants varied considerably between constructs (Table 2). 19,200 plaques were recovered from the co-transformed G1/G2 libraries, where only 3,480 total variants were expected in the pool. On the contrary, 460 plaques were observed for the F2 library, when 2,280 total variants were in the starting plasmid pool. The underrepresentation of the F2 library at this scale could be remedied by transforming more competent cells. We expect that differences between fragments in the number of recovered viruses is due to in part to the tolerability of fragments to mutation, as inviable mutants would not form plaques. Estimates from deep mutational scanning of model proteins show that approximately 18-38% of all mutations are nonviable (Firnberg et al. 2014, Faber et al. 2019), and this percentage may be larger for structural proteins like F and G.

**Table 2.**
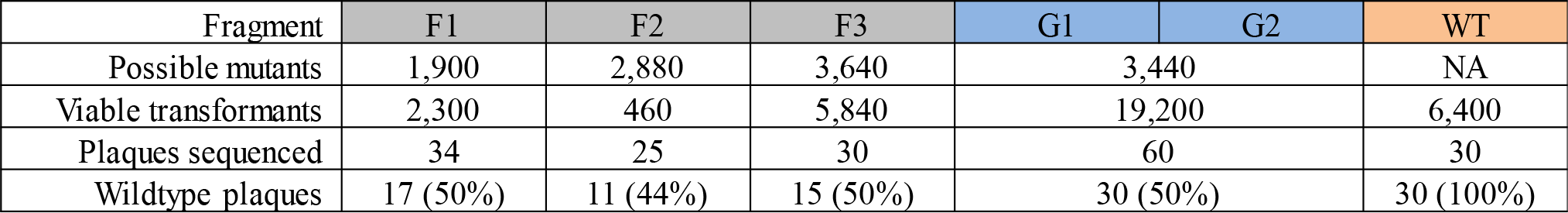
Characterization of viable virus genotypes in mutant phage library. Summary table of number of possible viruses, number of viable viruses recovered from one transformation, and the number of mutant and wildtype genotypes recovered. No mutant genotypes were observed more than once.

To test the validity of the mutagenesis and phage reconstitution method, we picked about 30 plaques generated from each library to Sanger sequence. Of these, 40-50% were wildtype and the remaining contained exactly one mutation (Table 2). No variants were observed more than once, supporting the deep sequencing results showing that the libraries consist of thousands of variants. Interestingly, none of the exact substitutions in these sequences have been previously observed (Supplementary Table S6), although mutations at the same residues (G106, F29, F45, F84, F185, F204, F205, and F393) have been seen in previous experiments (Wichman and Brown 2010). In all cases the percentage of WT plaques is slightly higher than in the starting plasmid pools (p-values F1: 0.001, F2: 0.088, F3: 0.015, G: 0.001 1-tailed binomial distribution), consistent with purifying selection against mutations (Table 1). In contrast, for the control where all plasmids in the Golden Gate reaction were WT, 0% (0/30) plaques contained mutations in the encoded regions.

## Discussion

Here we present a novel method for generating comprehensive single-site and multi-site saturation mutant libraries of the spike and capsid proteins of the bacteriophage ΦX174. This method uses unamplified oligo pools, nicking scanning mutagenesis, and Golden Gate cloning of the fragmented genome.

The fragment-based system provides an ideal way to generate libraries for the study of epistasis and multi-mutational step regions of the neighborhood around wild type. When a gene is in multiple fragments, the number of mutational variants we expect from the assembled mutant fragments is the product of their individual variant number. Hence, since G1 contains 2640 variants and G2 contains 800 variants, they can be combined to generate 2.1 million double mutation variants. Further, nothing requires the fragments to be from the same gene. For example, since the F and G proteins bind to each other to form the capsid of phiX174 (Fig. 2), libraries combining mutations in F with mutations in G will capture mutations that can readily interact in the protein and have the potential to generate interesting instances of epistasis (Fig 2). The central challenge then becomes how to survey the vast size of these combinatorial plasmid libraries. Right now, our transformational efficiencies vary, but even at the upper end of hundreds of thousands of transformants per reaction, only a fraction of the total diversity in the library can be surveyed.

**Figure 2.**
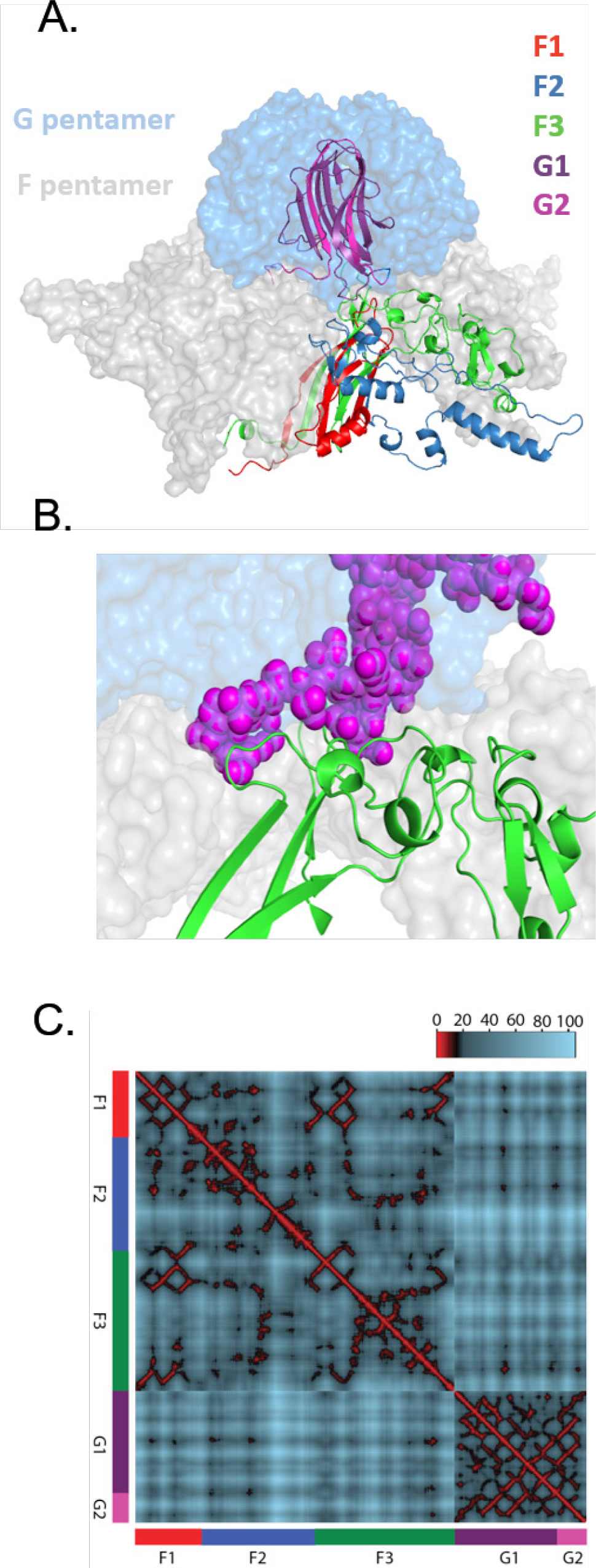
Structural view of potential epistatic interactions between F and G libraries. **A.** Pentameric complex between F and G shown in surface view. For both F and G, one monomer is shown as a ribbon color-coded according to the saturation mutagenesis library prepared in this work. **B.** Close-up of interaction surface between the F3 sub-library of F protein (green ribbons) and G2 sub-library of G protein (purple spheres). **C.** Distance mapping between alpha carbons in F and G crystal structure. Residues within 10 angstrom are colors red, 10-20 are black, and >20 are blue. Euclidian distances calculated from F-G crystal structure. Heatmap shows that many opportunities exist for close proximity (interactions) both within and between fragments.

We are fortunate that our model virus has a small genome, under 6,000 nts, which enabled us to assemble the 14 non-toxic fragments with sufficient transformants for near-complete coverage of the user-defined mutations. This method can be applied with little modification to other phages with small genomes like the microviridae. However, most biotech-relevant phages have larger genomes, and extension of this method to other viruses would encounter the following technical challenges. First, increasing genome sizes decrease transformational efficiencies, and larger genomes have increased susceptibility to DNA shearing (Rogers and Bendich 1988). Both size-dependent effects can disrupt the integrity and coverage of the phage libraries (Edwards and Rohwer 2005). Second, our method includes restrictions on the DNA sequences that can be used. Golden Gate cloning requires a genome without unique Type IIS restriction sequences, while nicking mutagenesis requires that there be only one orientation of the BbvCI nicking site within the template DNA fragment. Third, ligating 14 fragments is close to the upper limit of Golden Gate cloning. Additionally, increasing the size of individual fragments is difficult as the presented method depends on replicating plasmids in *E. coli*, and larger fragments are more likely to be toxic in vivo.

Based on the above considerations, adapting this method to larger viruses will require modifications for the genome assembly steps. There are a variety of methods available for attempting to improve genome library assembly and amplification. For improving genome assembly, we speculate that a combination of hierarchical assembly (Lee et al. 2015) with other yeast-based assembly methods (Ando et al. 2015) will allow for large viral genomes to be efficiently assembled. Yehl et al. 2019 build large libraries at targeted epitope regions by combining methods; plasmid-based mutagenesis and cloning followed by recombination with wild-type phage during infection. Finally, we speculate that the Tx.Tl cell free expression system (Garamella et al. 2016) will be able to produce the viral libraries with less bias than in vivo amplification in *E. coli*. In summary, we have presented a method for efficient deep mutational scanning of the bacteriophage ΦX174. We anticipate that this method will find utility in fundamental molecular evolution studies as well as translate to potential medicinal applications.

## Methods

### Reagents

All purchased enzymes and DNA purification kits were from New England Biolabs, antibiotics were purchased from GoldBio, other chemicals were purchased from Sigma-Aldrich. Individual primers were purchased from Integrated DNA Technologies.

### Segmentation of the ΦX174 genome

A phage assembly platform for ΦX174 was devised following (Donaldson et al. 2008). The ΦX174 chromosome was divided into 14 genomic fragments designed to avoid host cell toxicity by separating genes from their promoters and breaking large genes into multiple segments (Supplemental Note S1). Each segment is flanked by unique five nucleotide overlaps of WT ΦX174 sequence so that they can be amplified from the ancestral ΦX174 using PCR primers designed to incorporate terminal BsmB1 restriction sites. Amplicons were cloned into pCR2.1 using the Invitrogen TOPO TA cloning system (Life Technologies, Grand Island, NY) and verified by Sanger sequencing.

### Introduction of nicking site

The gene fragments – F1, F2, G1, G2 – in the pCR2.1-TOPO plasmid (Supplemental Notes S1 and S2) had the BbvCI nicking site introduced via overhang PCR, type I restriction enzyme cutting, and ligation. First, PCR was performed with overhang primers (Supplemental Table S1) to introduce the BbvCI site. Standard Phusion Polymerase HF reaction conditions were used with 4 ng of template DNA, cycling conditions as follows: 98°C - 1 min, 25× cycles of: 98°C – 10 seconds, 67°C 15 seconds, 72°C 2.5 minutes, followed by 72°C for 10 minutes. PCR products were run on a 1% agarose gel stained with SYBR™ safe stain (Invitrogen) and the DNA bands at ~4600 bp were extracted using a Monarch^®^ DNA Gel Extraction Kit. Next, 1 μg of the PCR product was digested with KpnI (20 U) in NEB Buffer 1.1 at 37°C for two hours. Digested DNA was cleaned and concentrated with a Monarch^®^ PCR and DNA Clean and Concentrate Kit and eluted into 20 μL nuclease free H2O. One microliter of the purified and digested DNA was ligated using T4 DNA ligase at ~25°C for 1 hour in standard conditions in a 20 μL reaction. Five microliters of the ligation reaction were transformed into chemically competent XL1-Blue *E. coli* via standard protocols. Cells were plated on LB agar containing 100 μg/mL carbenicillin and 50 μg/ mL kanamycin and grown at 37°C for ~16 hours. Cells were picked from transformation plates and grown in 50 mL TB with 100 μg/mL carbenicillin and 50 μg/mL kanamycin for ~12 hours, cells were pelleted, and DNA purified using compact midi-preps.

### Construction of single site mutant libraries

Comprehensive mutant libraries were generated using nicking mutagenesis (NM) as in Wrenbeck et al. 2016 with modifications for using oligo pool mutagenic primers as noted in Medina et al. 2019. A single oligo pool encoding for all possible single missense and nonsense substitutions in F and G was designed using the custom python scripts from Medina et al. 2019 and custom synthesized by Agilent (full sequences of all oligos are given in the associated file “VMA_supplemental_data.csv”). Oligo pools were designed with 20-24 bases of gene overlap flanking the mutated codon. Codons were chosen based on *E. coli* codon usage frequency, where the highest frequency codon that encoded the mismatch mutation was selected. We printed 14,853 unique oligos, where 14,853 = 12,453 ΦX174 primers– 2,400 other primers. These oligos encode WT (593 primers) and all 20 possible single point mutations (including stop) (11,860 primers). This oligo pool was used directly in NM without further amplification using 2 μg of the relevant golden gate plasmid as a template. Libraries were electroporated into high efficiency electrocompetent XL1-Blue *E. coli* (Agilent cat #: 200228) using 1 mm electroporation cuvettes at 1200V using a Eppendorf Eporator. Cells were plated on large bioassay plates (245mm × 245mm × 25mm, Sigma-Aldrich) containing LB agar + 100 μg/mL carbenicillin and 50 μg/mL kanamycin and grown at 37°C for ~16 hours. Plates were scraped in 10 mL plain LB, broken into ~1.2 mL aliquots, pelleted, and stored −80°C. DNA was purified from 1× aliquot using a Monarch^®^ Plasmid Mini-Prep Kit.

### Illumina sequencing prep and analysis

Purified library DNA was prepared for deep sequencing as in Kowalsky et al. (2015) using the primers listed in Supplemental Table S2 and gene tiling as specified in Table 1. DNA was Illumina sequenced on a Mi-Seq platform with 250 BP paired end reads. The University of Colorado BioFrontiers Sequencing Core performed the Illumina sequencing. Data was processed using PACT (Klesmith and Hackel 2018) to determine the library coverage with the following changes to the default options in the configuration file: fast_filter_translate: qaverage = 20, qlimit = 0; enrichment: ref_count_threshold = 5, sel_ count_threshold = 0, strict_count_threshold = True. Library coverage statistics are shown in Supplemental Table S3 and S4.

### Assembly of mutant genomes

We pooled plasmid DNA containing all 14 of the phage DNA fragments (500 ng each) and digested them with BsmB1 (Fermentas Fast Digest, Life Technologies, Grand Island, NY) for 30 minutes to 1 hour at 37°C. The digested plasmids were subjected to agarose gel electrophoresis for 10 to 15 minutes using a 1.2% agarose gel to separate the vector from the inserts. The inserts were excised from the gel, purified using the GeneJET gel extraction kit (Fermentas), ligated overnight at 14°C with T4 DNA ligase (Promega Corporation, Madison, WI), and transformed by electroporation into 50 μl of XL1-Blue electocompetent cells. The transformation mix was resuspended with 950 μl of SOC and either plated immediately or incubated for up to 120 minutes (Supplemental Table S5) at 37°C in the presence of DNase1 (Supplemental Figure S10) to allow for cell lysis and the removal extracellular DNA, respectively. Aliquots of the cell lysates were added to 3 ml of ΦLB top agar containing 50 μl log-phage E. coli C cells and plated onto a ΦLB agar plate. After four to five hours of incubation at 37°C, recombinant phage plaques were visible and plates were removed from the incubator. To verify that the recombinant phage encoded the intended sequence, we picked about 30 plaques for each intended mutational target and Sanger sequenced the entire targeted gene. We also sequenced the F and G genes for about 30 wildtype plaques to assure that no mutations were naturally accumulating. Briefly, individual plaques were picked with sterile toothpicks and placed in 200 uL ΦLB and gently swirled. 1 uL of this mix was used to PCR amplify approximately ½ of the ΦX174 genome using ΦX-0F (5’-GAGTTTTATCGCTTCCATG-3’) and ΦX-2953R (5’-CCGCCAGCAATAGCACC-3’) primers. Internal sequencing primers ΦX-979F (5’-CGGCCCCTTACTTGAGG-3’) and ΦX-1500R (5’-TTGAGATGGCAGCAACGG-3’) were used to sequence gene F. ΦX-2953R was used to sequence gene G. PCR cleanups and sequencing was done at Eurofins Genomics. PCR reaction conditions were: 5 uL10X Taq buffer, 2.5 uL 10 uM ΦX-0F primer, 10 uM ΦX-2953F primer, 0.8 uL 12.5 uM dNTPs, 0.5 uL Taq polymerase (NEB #M0273), 1 uL template, 37.7 uL H20. Thermocycling conditions were 1 cycle at 95°C for 2 min, 30 cycles at 95°C for 15 sec, 52°C for 30 sec, 68°C for 2 min, 1 cycle at 68°C for 5min.

## Acknowledgements

We thank the members of the Whitehead, Wichman, and Miller labs for feedback and assistance in composing this work. We especially thank LuAnn Scott for experimental expertise on the ΦX174 system. This work was supported by the National Science Foundation BEACON Centre for the Study of Evolution in Action and an NSF EPSCoR T2 under award numbers DBI-0939454 and OIA-1736253 and the National Institute of General Medical Sciences of the National Institutes of Health under award numbers P20GM104420 and R01GM076040. The content is solely the responsibility of the authors and does not necessarily represent the official views of the National Institutes of Health.

## Data Availability

Raw sequencing reads for the truncated mutant libraries have been deposited in the SRA database: SAMN12660301-SAMN12660308. The processed data is available in the supporting information. Additionally, the oligo pool sequences, input files for generating the oligo pool, and an example of the command line input to execute the script from Medina-Cucurella et al. (2019) are available in the Supplemental Information associated with this paper.

## Supplemental Information

### Note S1: DNA sequences of ΦX174 viral capsid protein gene fragments

Sequences provided span from the 5’ EcoRV to the 3’ BamHI site (bolded sequences). Mutagenic regions are highlighted in pink, non-mutated coding regions are highlighted in blue, BsmBI sites are highlighted in green. Regions with overlapping protein coding regions on different reading frames are colored dark blue.

#### >Fragment_F1

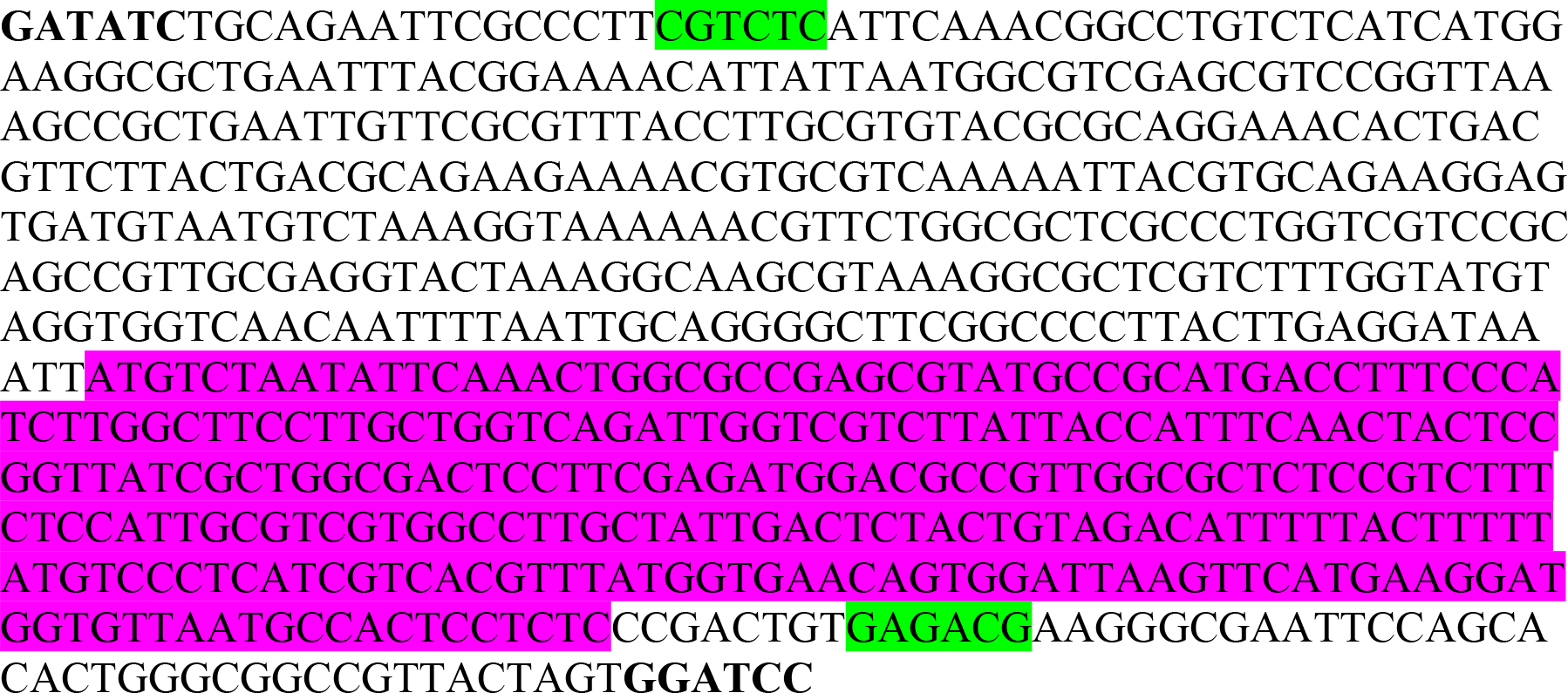

#### >Fragment_F2

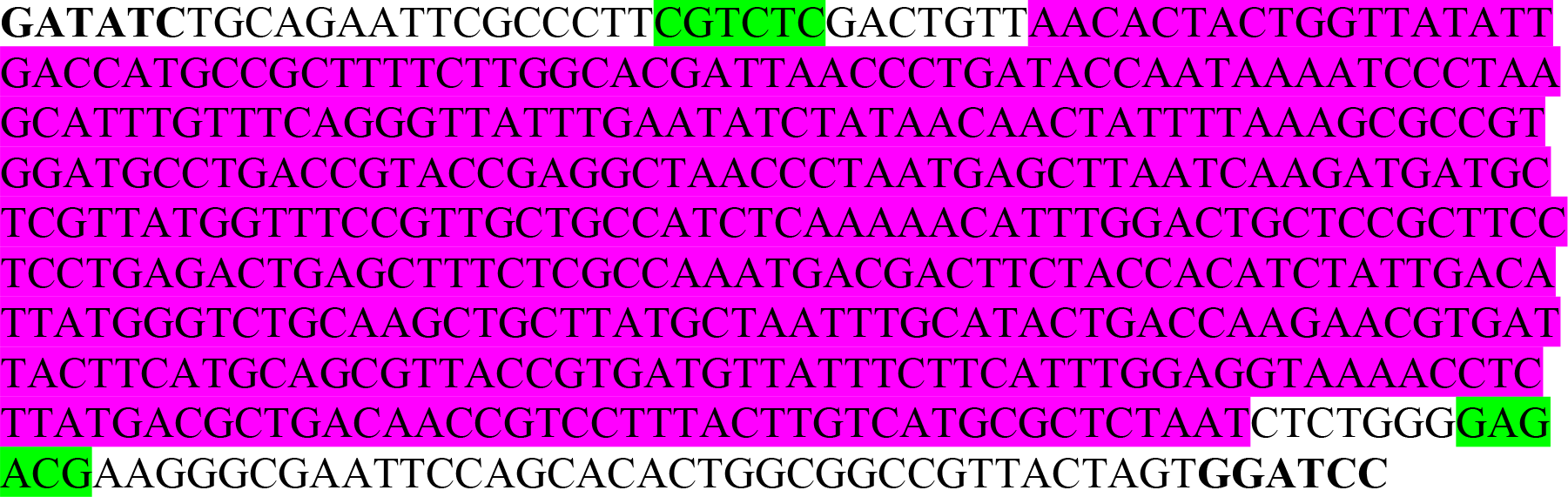

#### >Fragment_F3

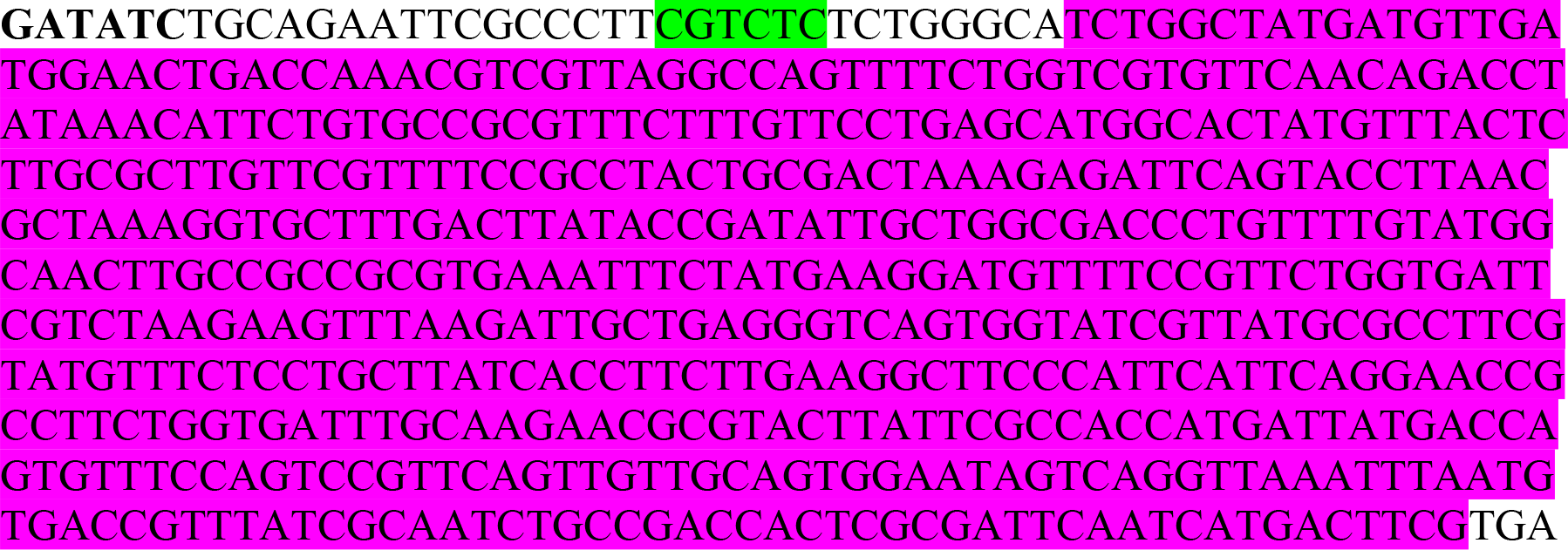
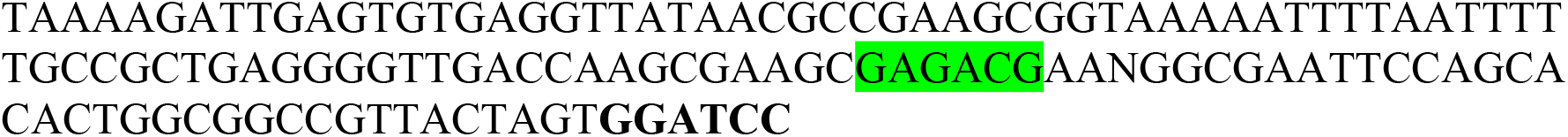

#### >Fragment_G1

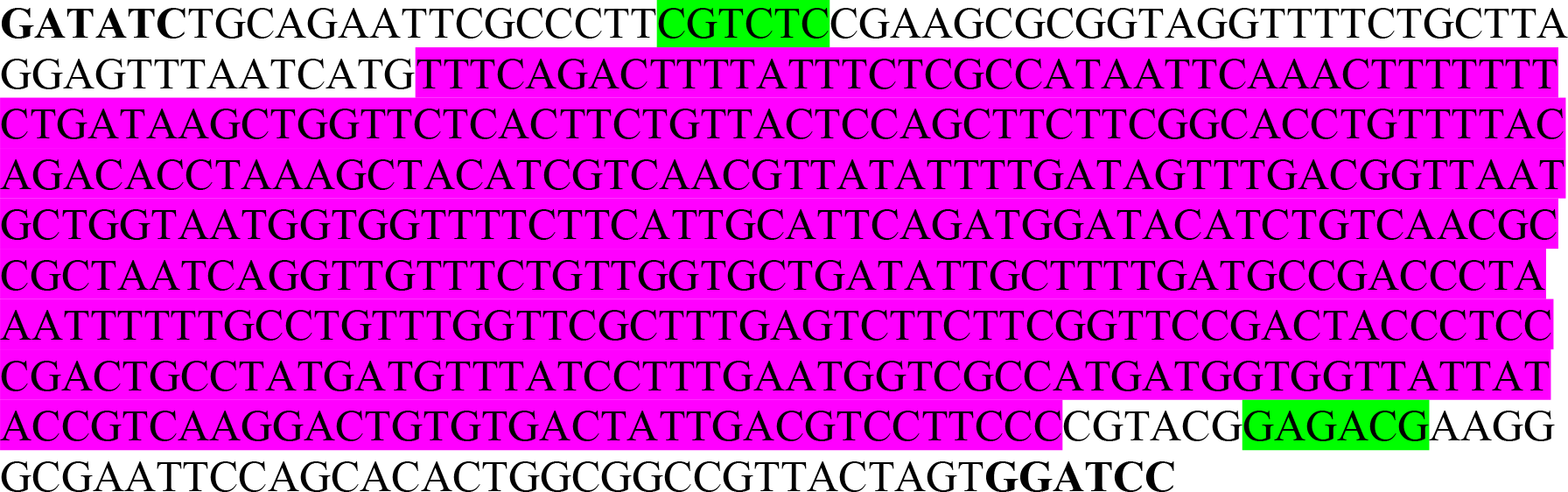

#### >Fragment_G2/H1

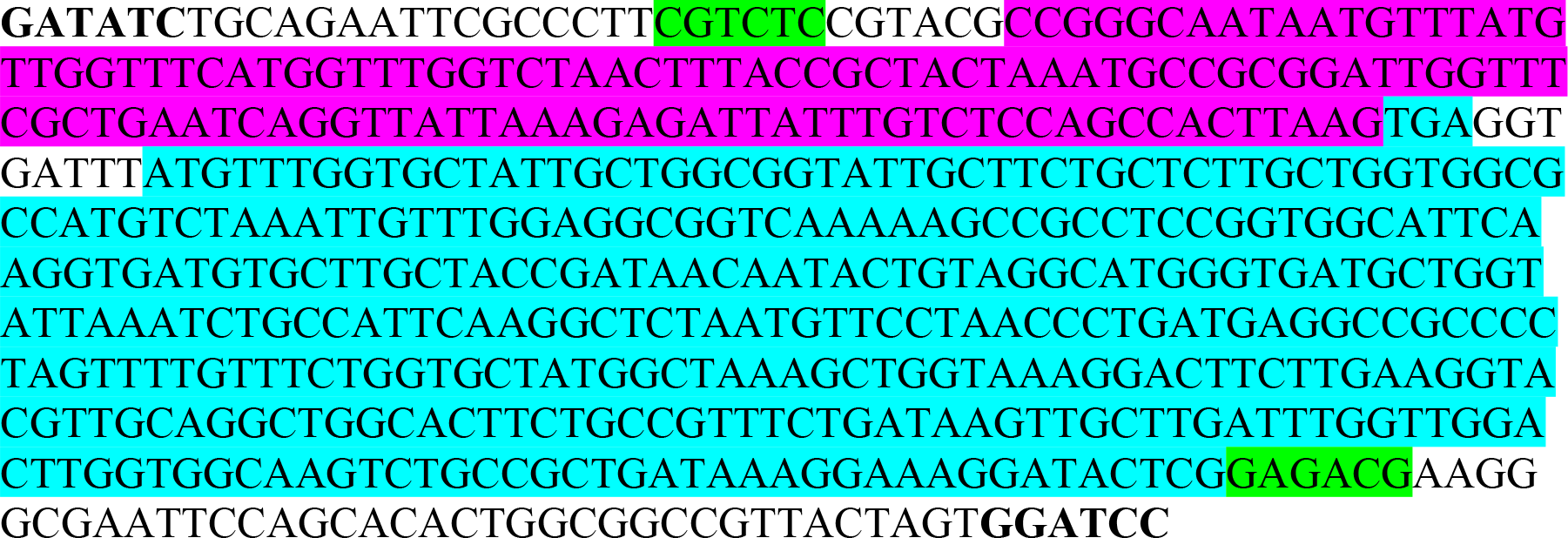

#### >Fragment_H2

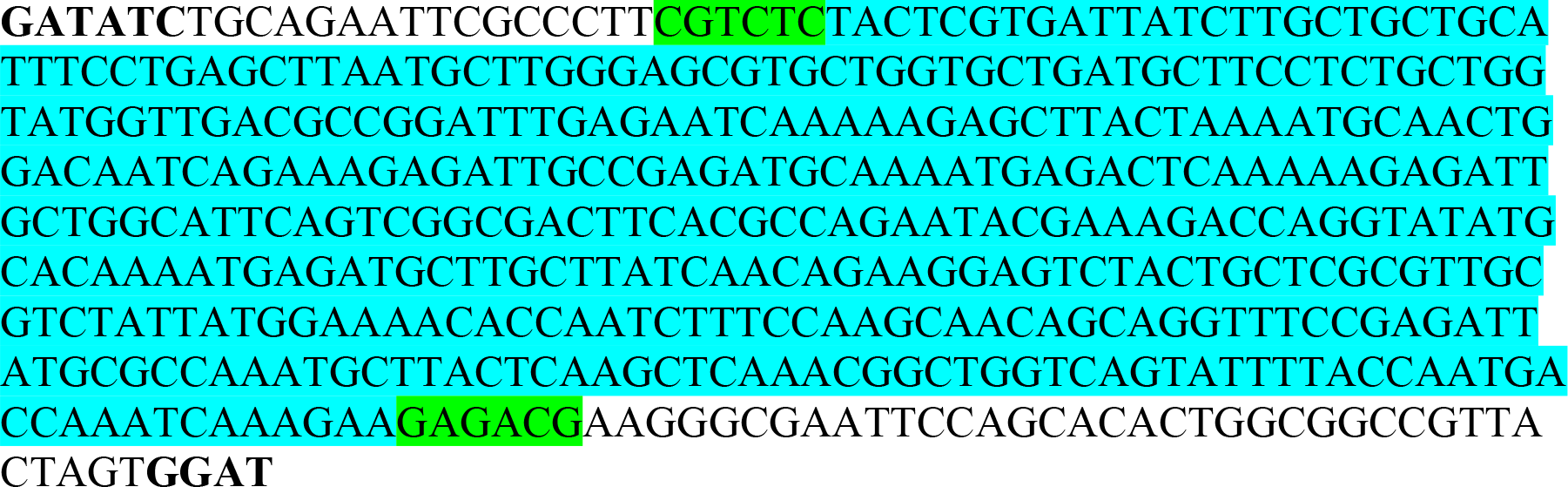

#### >Fragment_H3

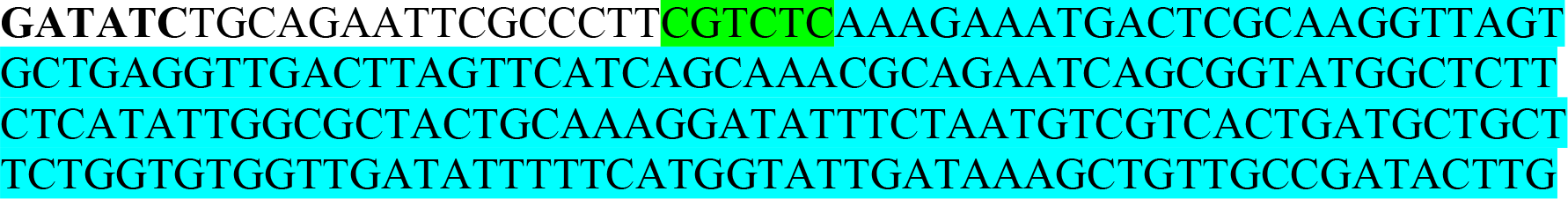
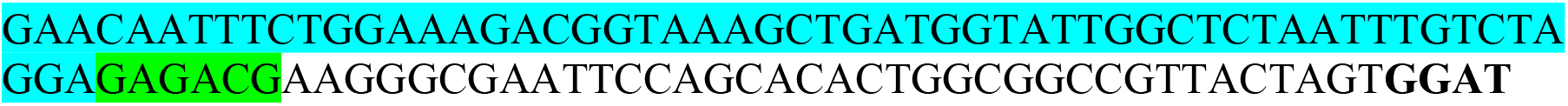

#### >Fragment_A1/H

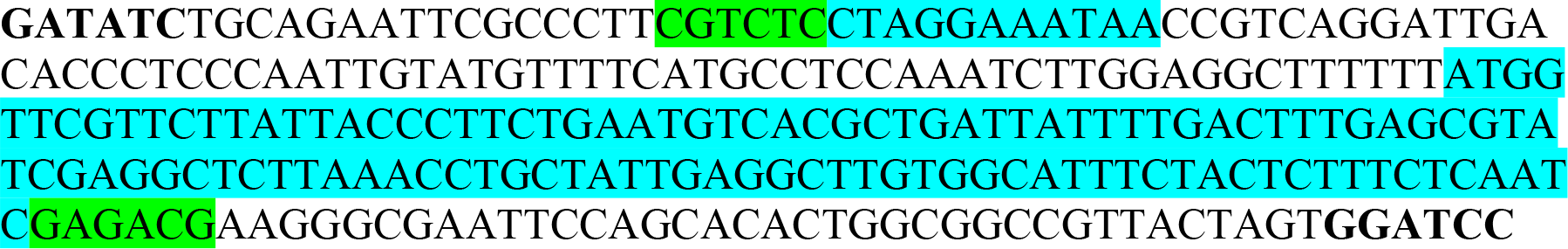

#### >Fragment_A2

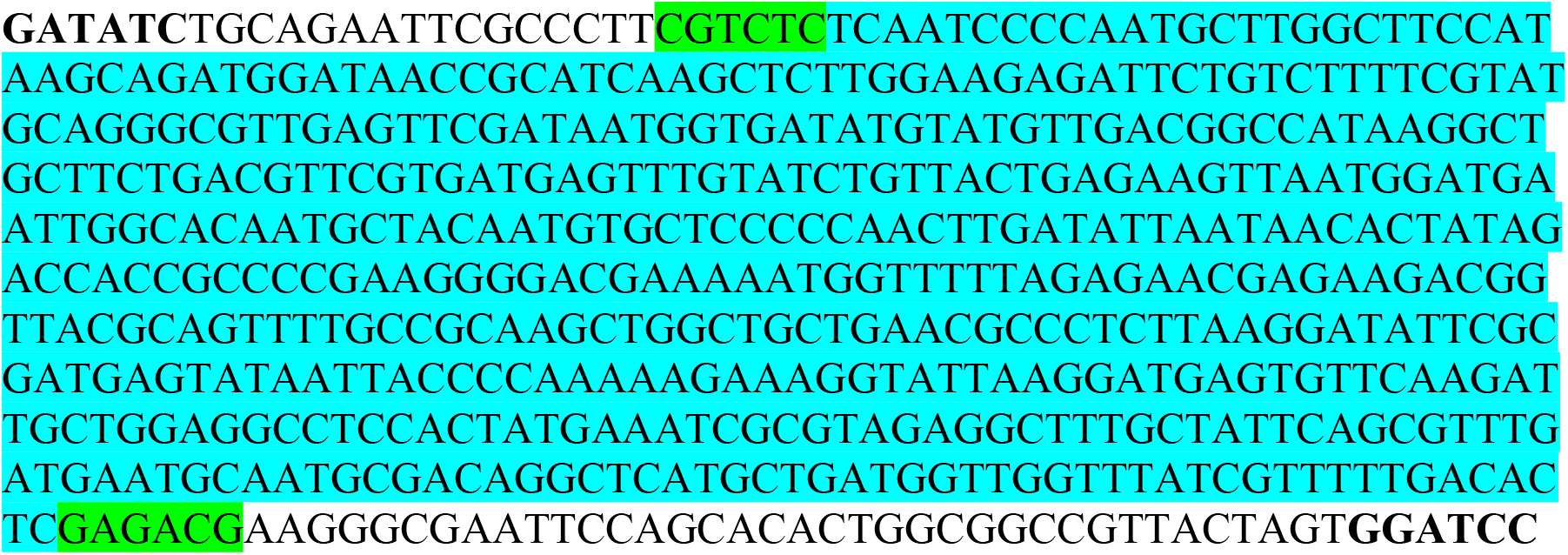

#### >Fragment_A3

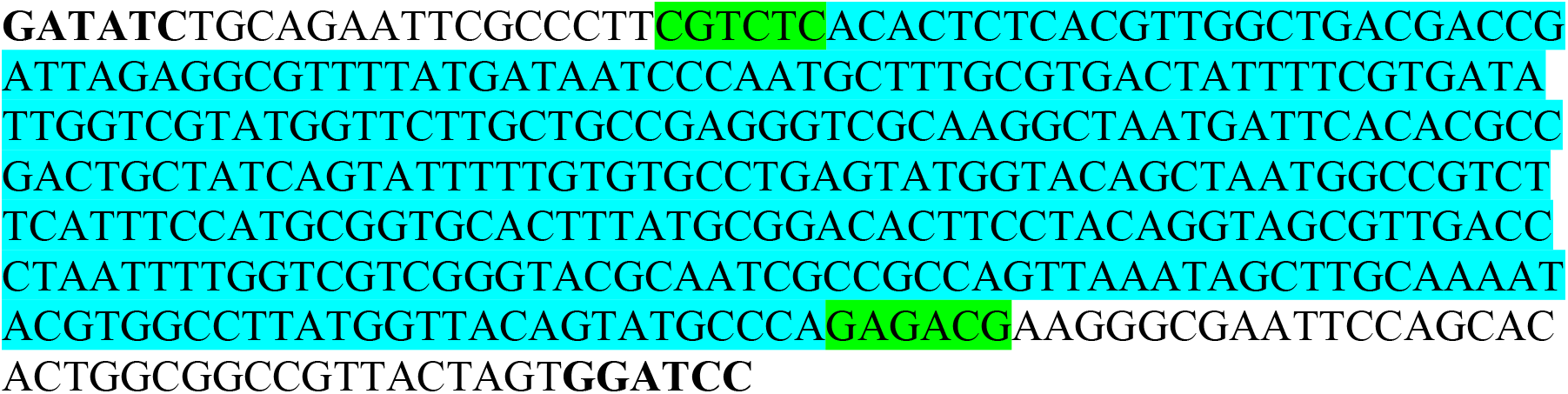

#### >Fragment_A4/B

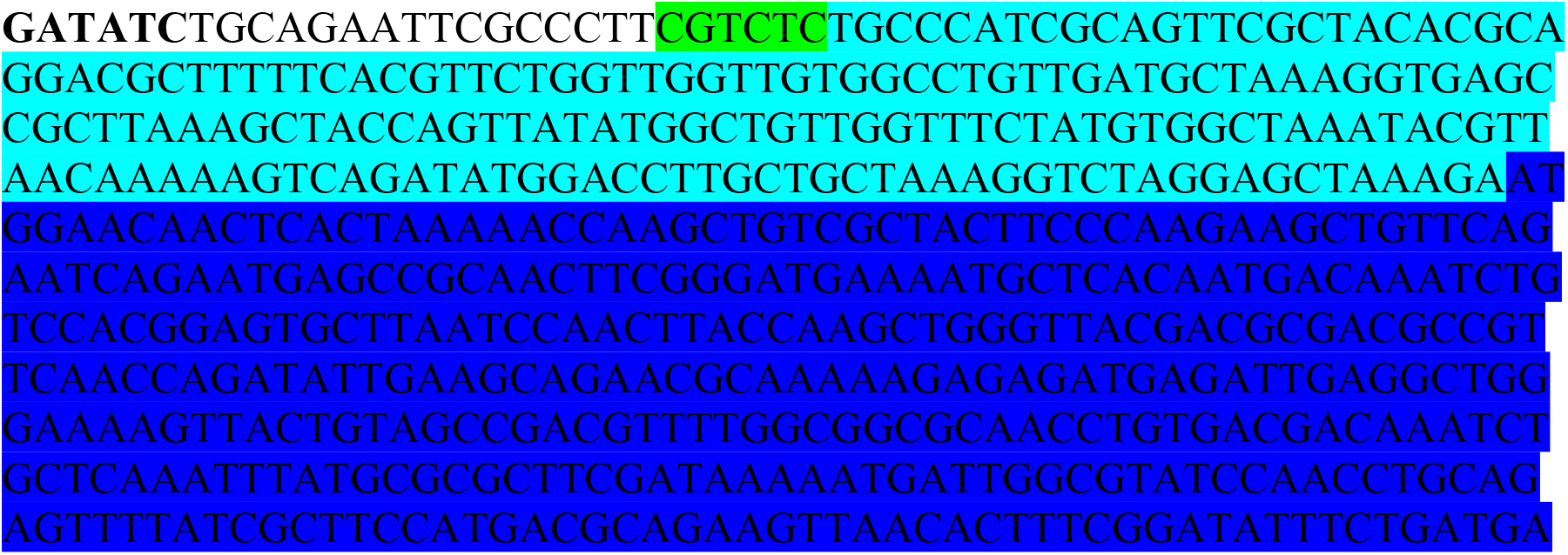
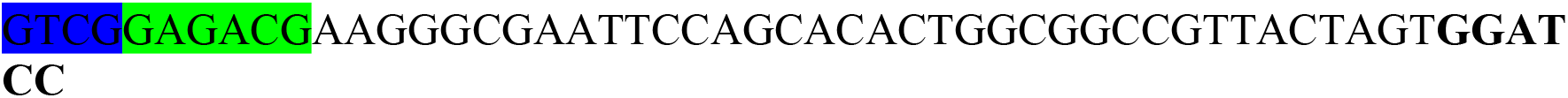

#### >Fragment_K/C/A

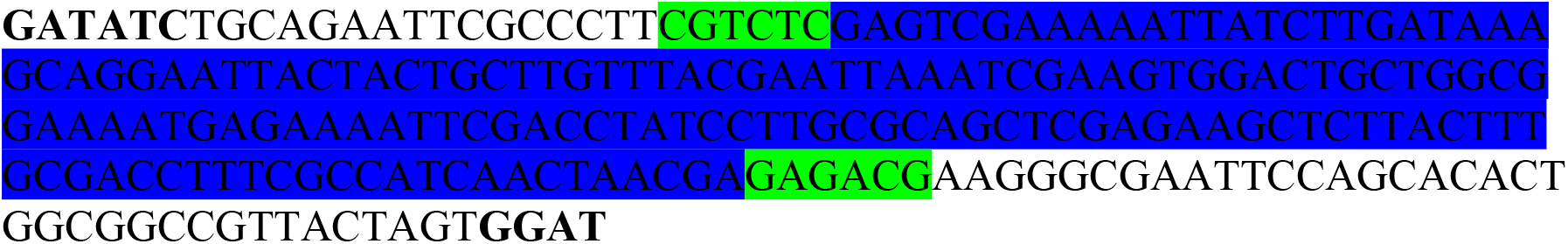

#### >Fragment_C/K

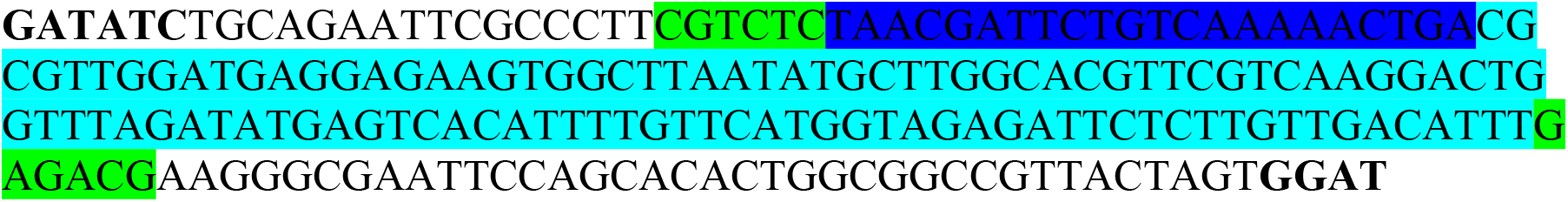

#### >Fragment_D/C/E

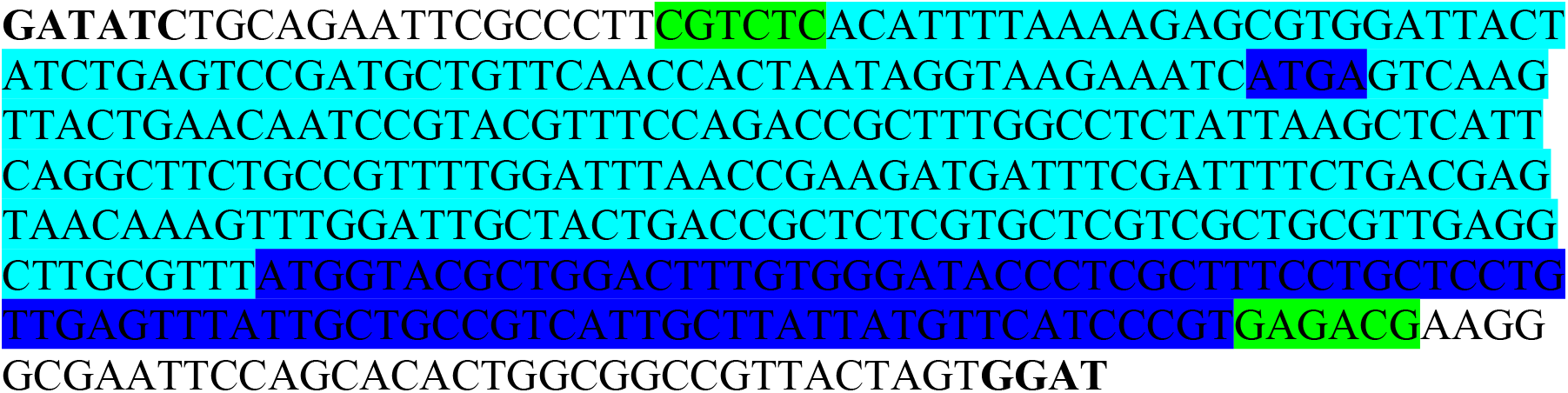

### Note S2: Amino acid sequences of ΦX174 mutagenized viral capsid protein gene fragments

Only mutagenized regions of gene fragments shown.

#### >Fragment_F1

MSNIQTGAERMPHDLSHLGFLAGQIGRLITISTTPVIAGDSFEMDAVGALRLSPLR

RGLAIDSTVDIFTFYVPHRHVYGEQWIKFMKDGVNATPL

#### >Fragment_F2

NTTGYIDHAAFLGTINPDTNKIPKHLFQGYLNIYNNYFKAPWMPDRTEANPNELN

QDDARYGFRCCHLKNIWTAPLPPETELSRQMTTSTTSIDIMGLQAAYANLHTDQE

RDYFMQRYRDVISSFGGKTSYDADNRPLLVMRSN

#### >Fragment_F3

SGYDVDGTDQTSLGQFSGRVQQTYKHSVPRFFVPEHGTMFTLALVRFPPTATKEI

QYLNAKGALTYTDIAGDPVLYGNLPPREISMKDVFRSGDSSKKFKIAEGQWYRY

APSYVSPAYHLLEGFPFIQEPPSGDLQERVLIRHHDYDQCFQSVQLLQWNSQVKF

NVTVYRNLPTTRDSIMTS

#### >Fragment_G1

FQTFISRHNSNFFSDKLVLTSVTPASSAPVLQTPKATSSTLYFDSLTVNAGNGGFL

HCIQMDTSVNAANQVVSVGADIAFDADPKFFACLVRFESSSVPTTLPTAYDVYPL

NGRHDGGYYTVKDCVTIDVLP

#### >Fragment_G2

PGNNVYVGFMVWSNFTATKCRGLVSLNQVIKEIICLQPLK

**Supplemental Figure S1.**
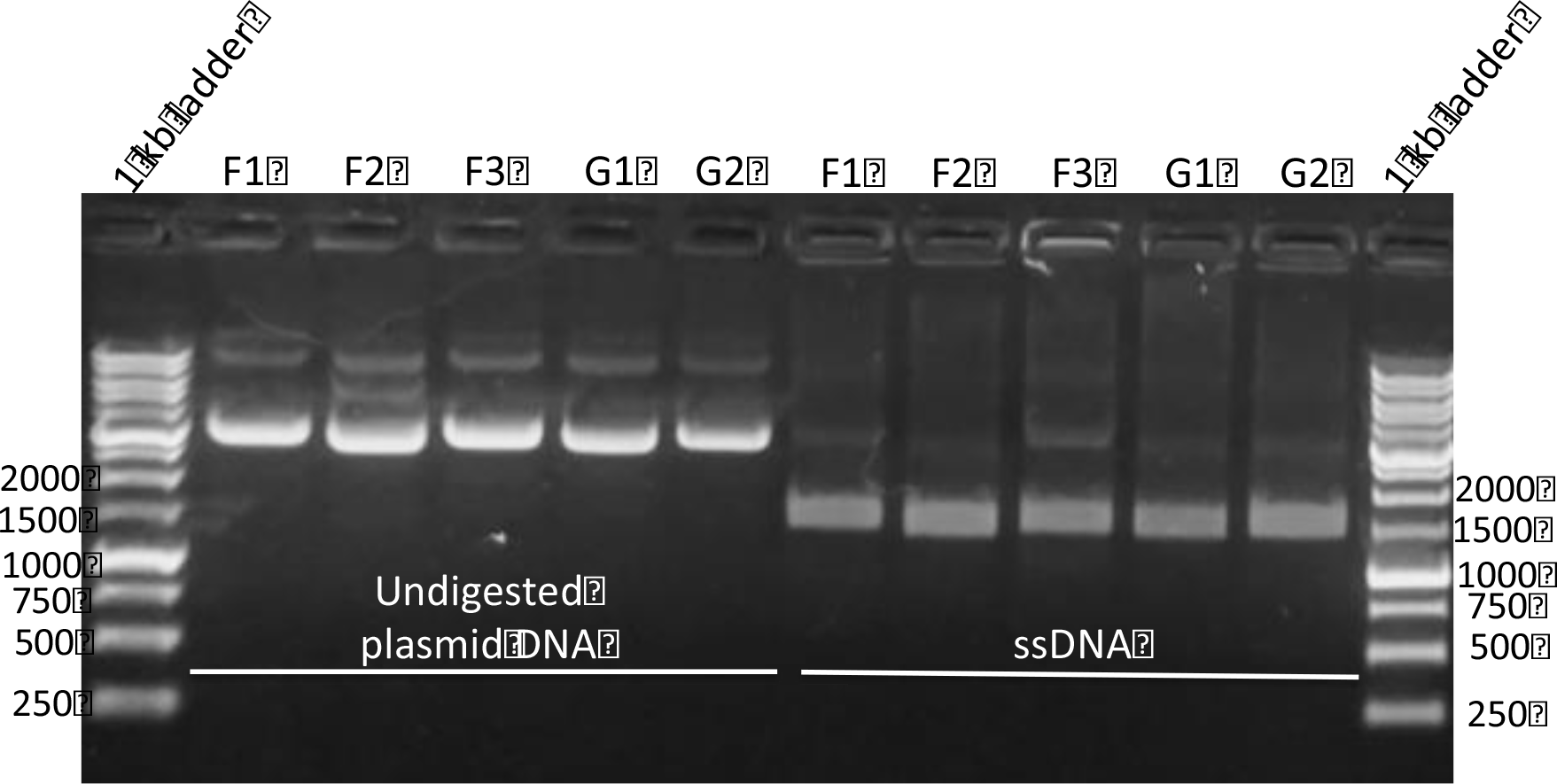
Introduction of nicking sites into shuttle vectors containing viral genes. Verification of the introduction of the BbvCI nicking site into the shuttle vector containing the viral genes is shown by the template prep step in the nicking mutagenesis protocol (generation of ssDNA). Samples were run on a 1% agarose gel with SYBR™ Safe DNA gel stain (Invitrogen) added before casting, the ladder used is the 1 kb DNA ladder from GoldBio.

**Supplementary Figure S2:**
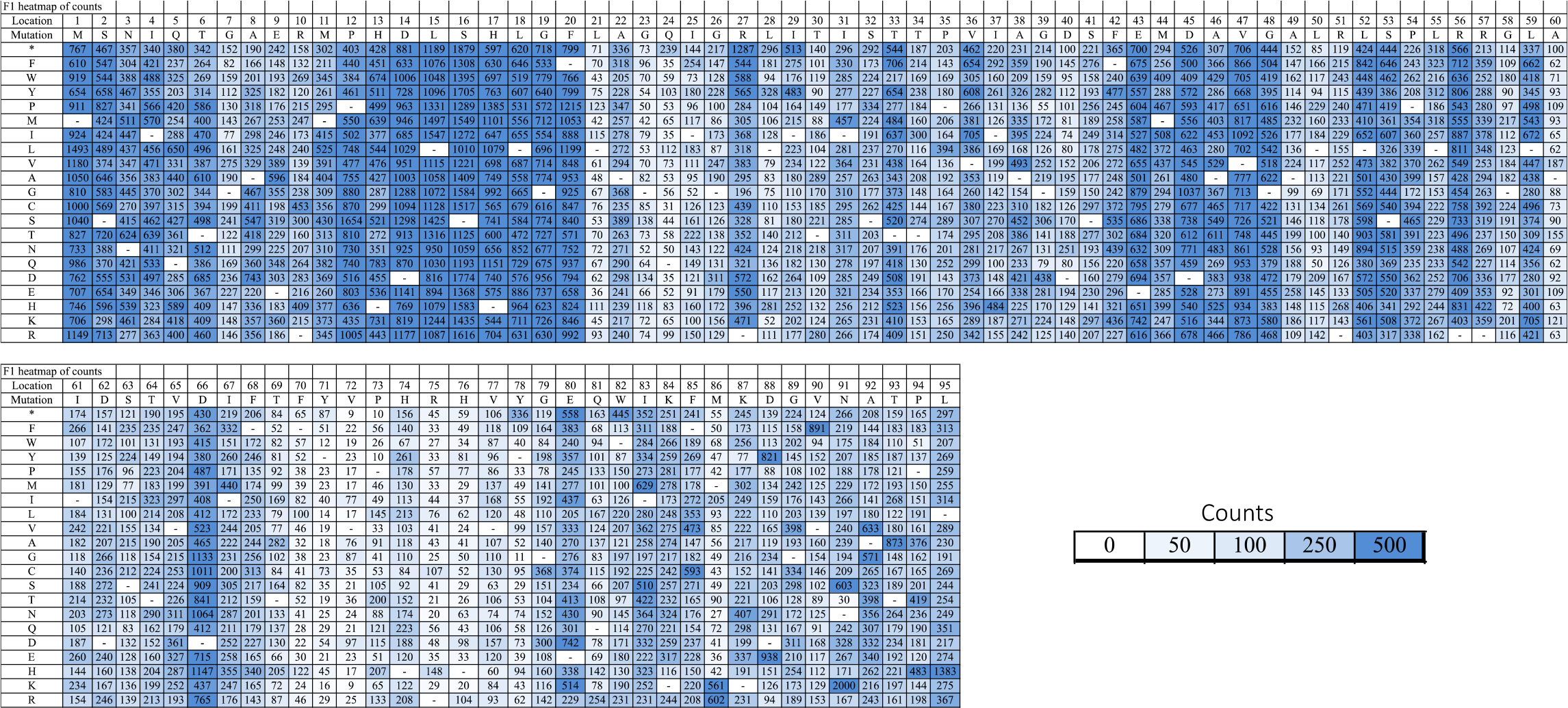
F1 heatmap of mutational frequency represented by number of sequencing read counts “counts”.

**Supplementary Figure S3:**
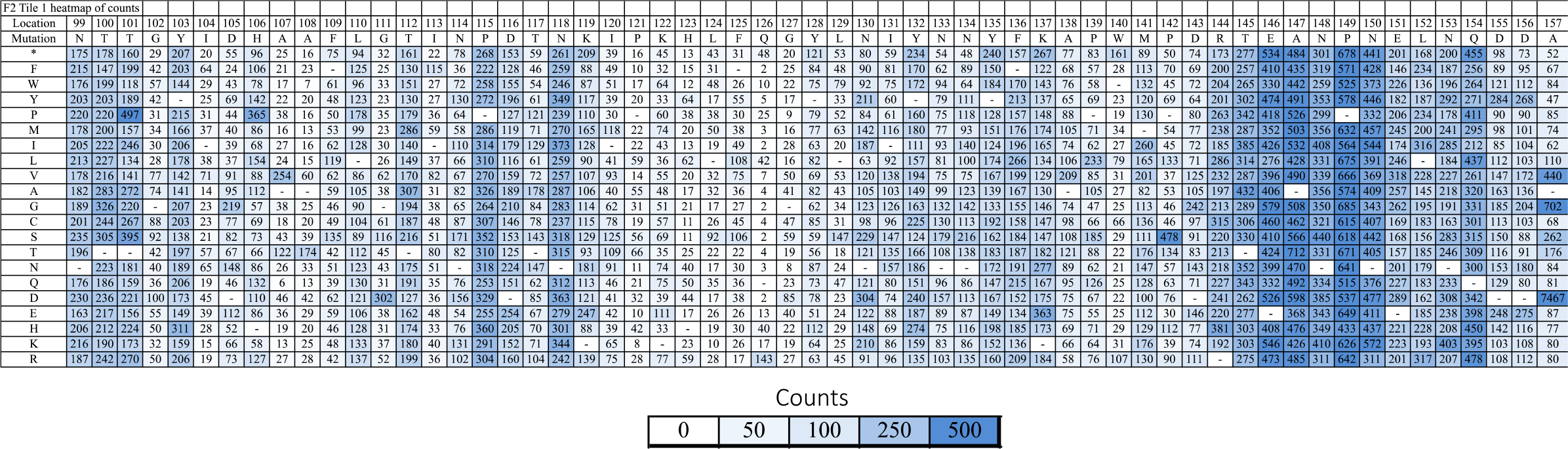
F2 tile 1 heatmap of mutational frequency represented by number of sequencing read counts “counts”.

**Supplementary Figure S4:**
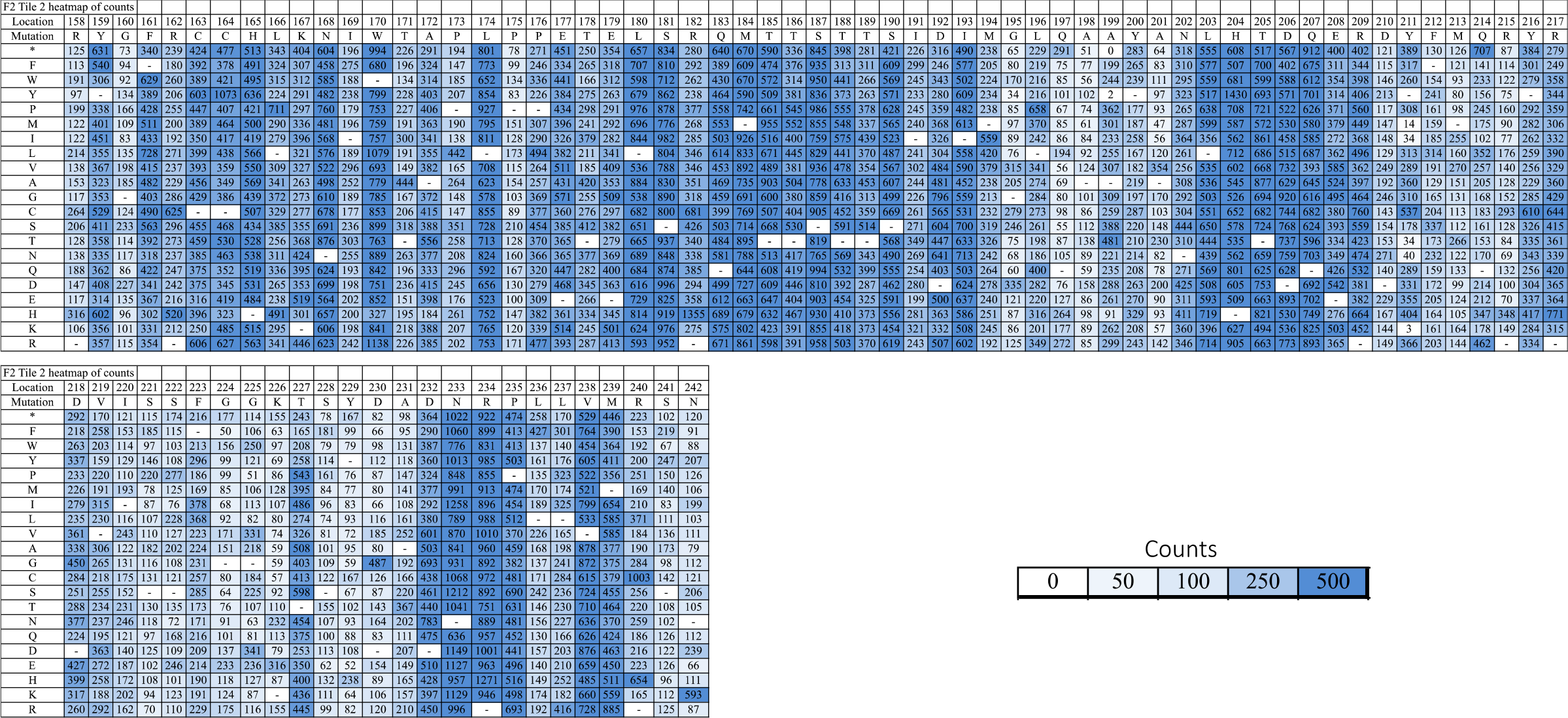
F2 tile 2 heatmap of mutational frequency represented by number of sequencing read counts “counts”.

**Supplementary Figure S5:**
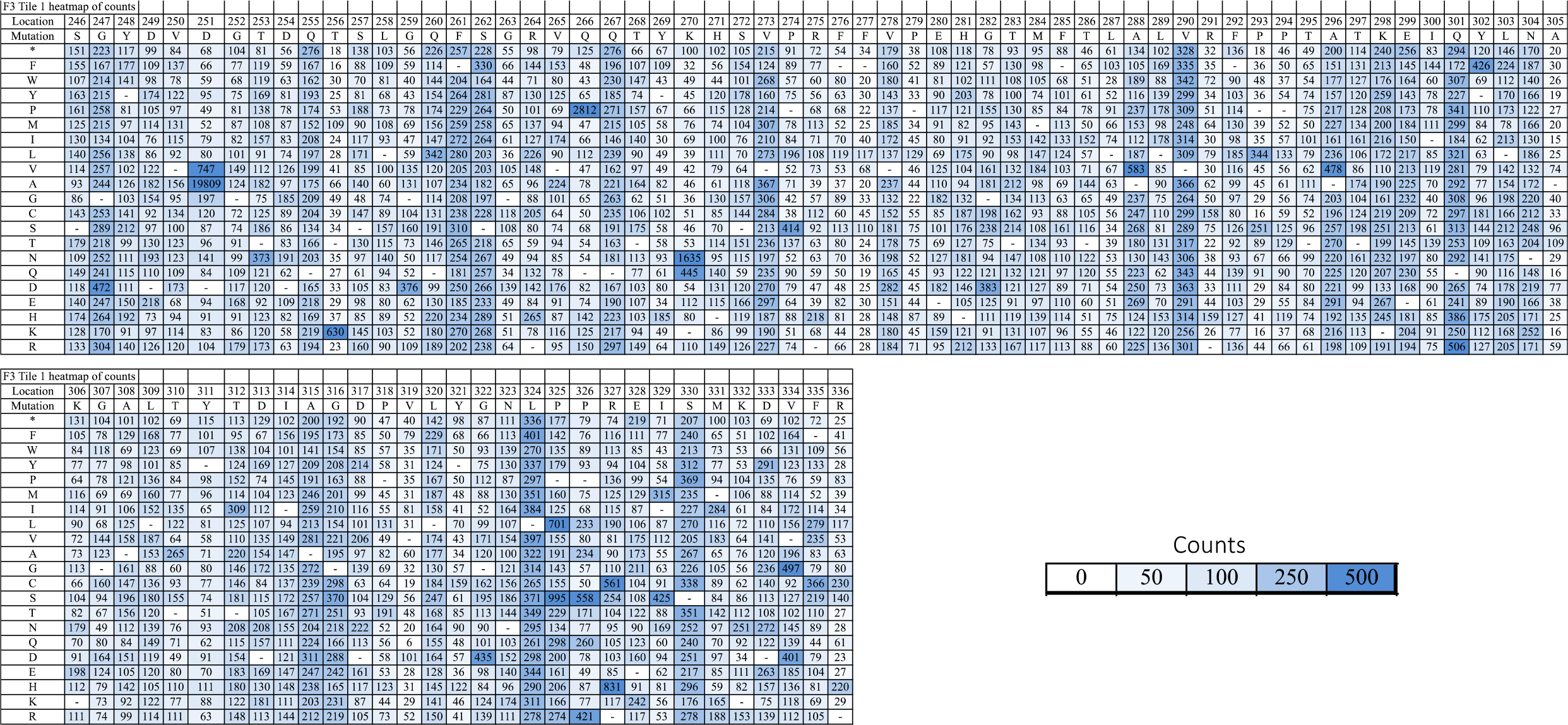
F3 tile 1 heatmap of mutational frequency represented by number of sequencing read counts “counts”.

**Supplementary Figure S6:**
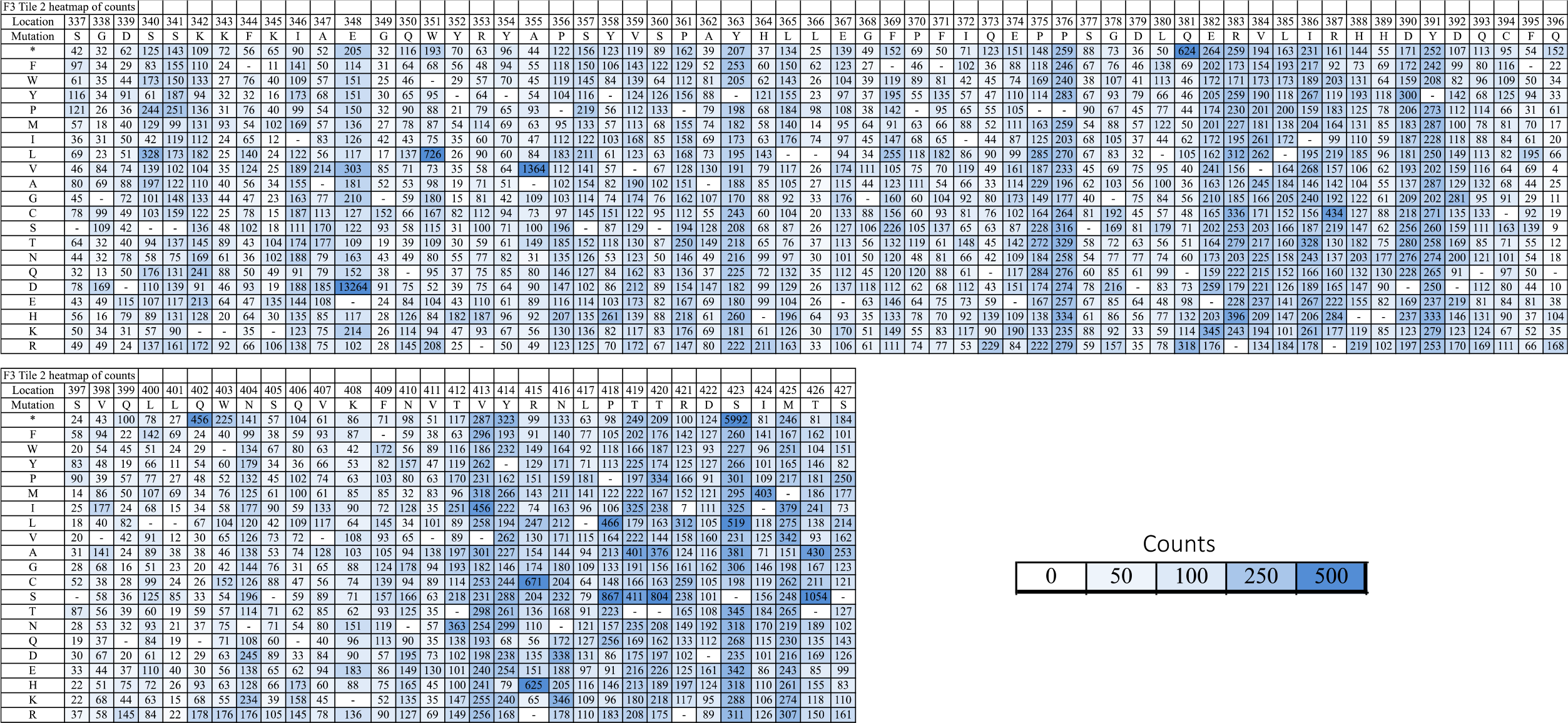
F3 tile 2 heatmap of mutational frequency represented by number of sequencing read counts “counts”.

**Supplementary Figure S7:**
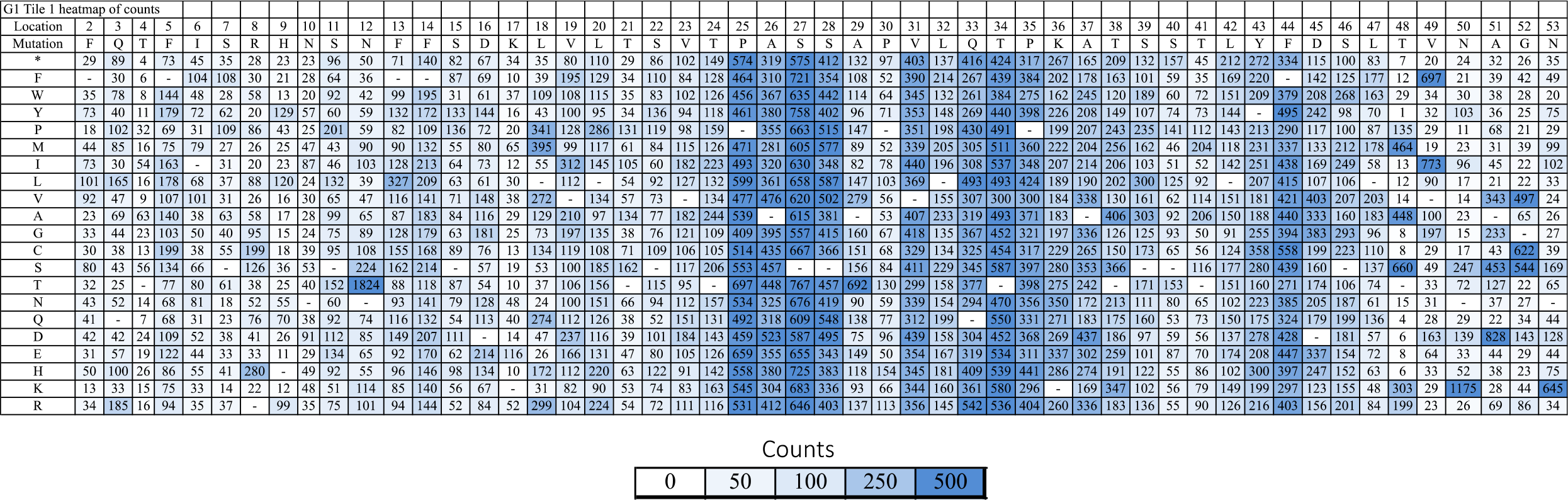
G1 tile 1 heatmap of mutational frequency represented by number of sequencing read counts “counts”.

**Supplementary Figure S8:**
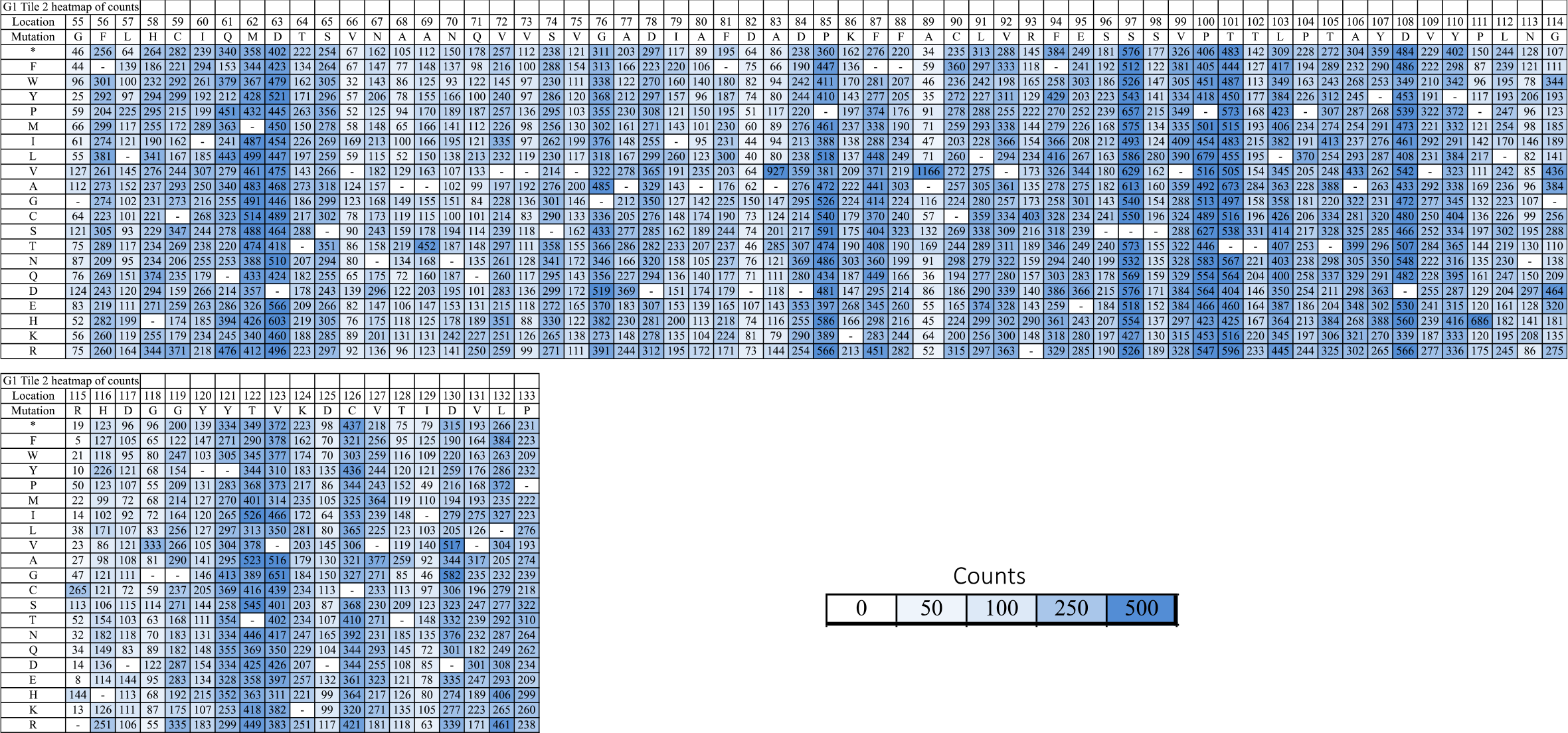
G1 tile 2 heatmap of mutational frequency represented by number of sequencing read counts “counts”.

**Supplementary Figure S9:**
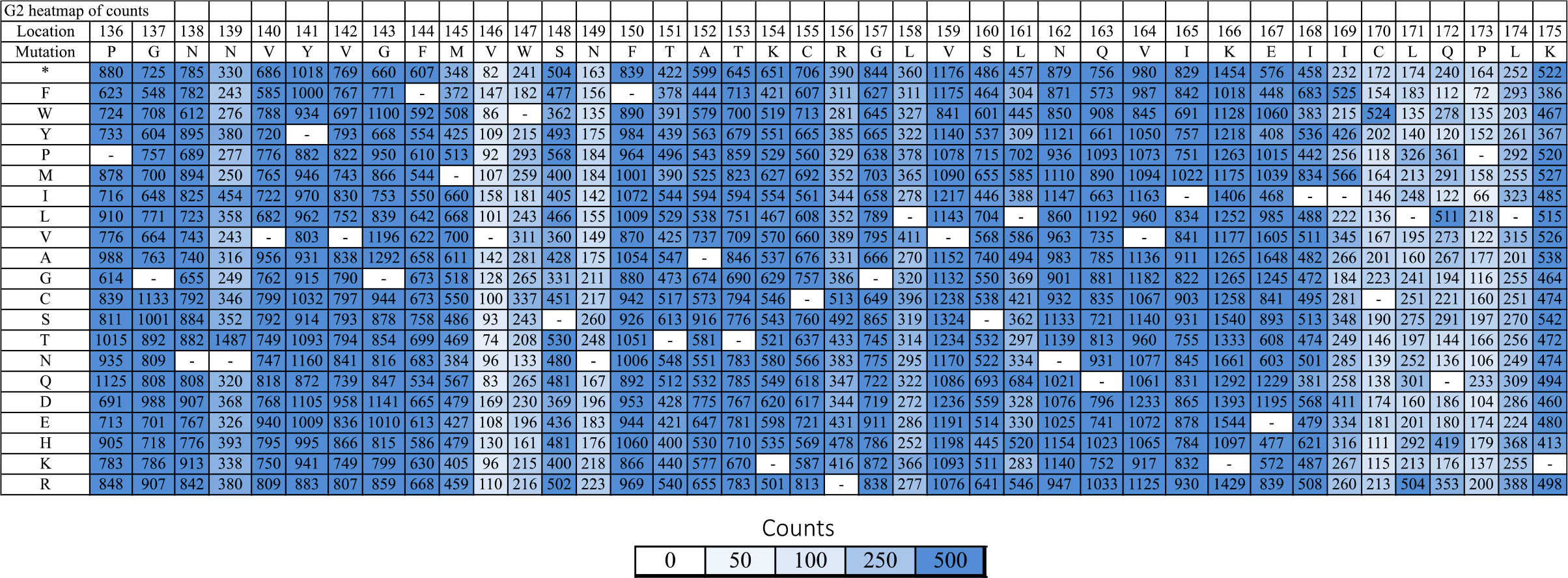
G2 heatmap of mutational frequency represented by number of sequencing read counts “counts”.

**Supplementary Figure S10:**
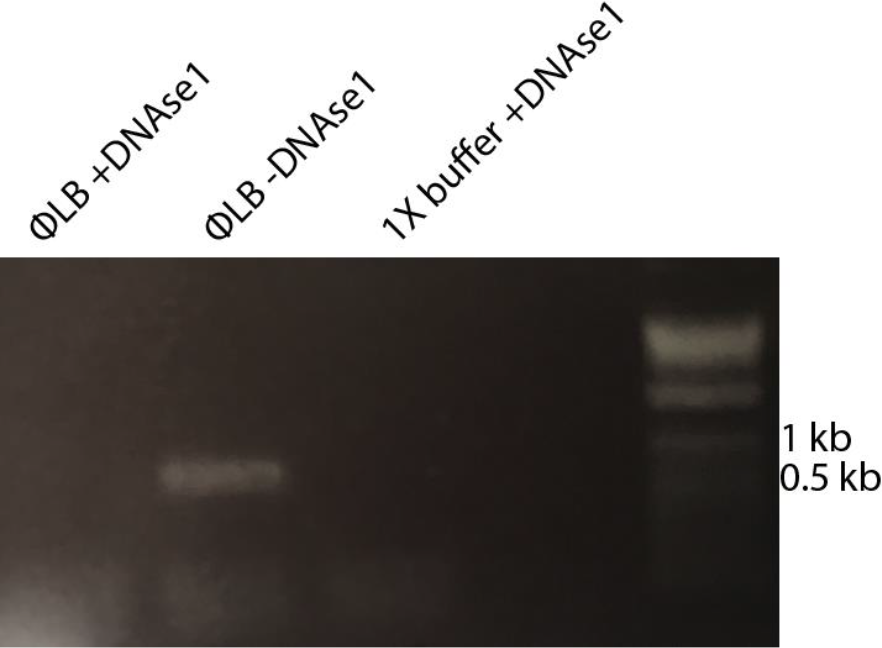
Agarose gel image showing effectiveness of DNase1 in ΦLB. 25 cycles of PCR were performed after a 1-hour 37°C incubation of WT ligation mix with DNase1. Primer sequences for phix_2375F and phix_2949R were 5’-ACACTGACGACATGGTTCTACAtctgcttaggagtttaatc-3’ and 5’-TACGGTAGCAGAGACTTGGTCTgcaccaaacataaatcacc-3’. These primers include annealing sequences for Illumina 2^nd^ round barcoding primers.

**Supplemental table S1.**
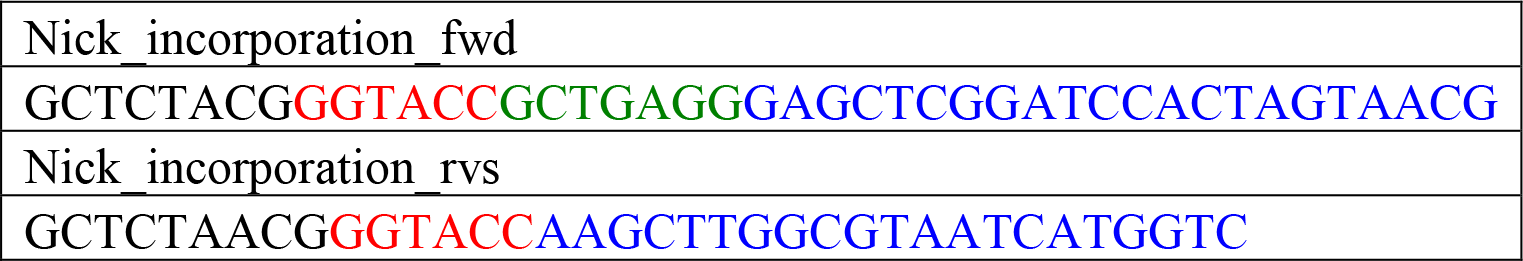
Primers for incorporating BbvCI nicking sites into the pCR2.1-topo shuttle vector. Blue text is the overlap region with the shuttle vector, red is the KpnI site, green is the BbvCI site.

**Supplemental table S2.**
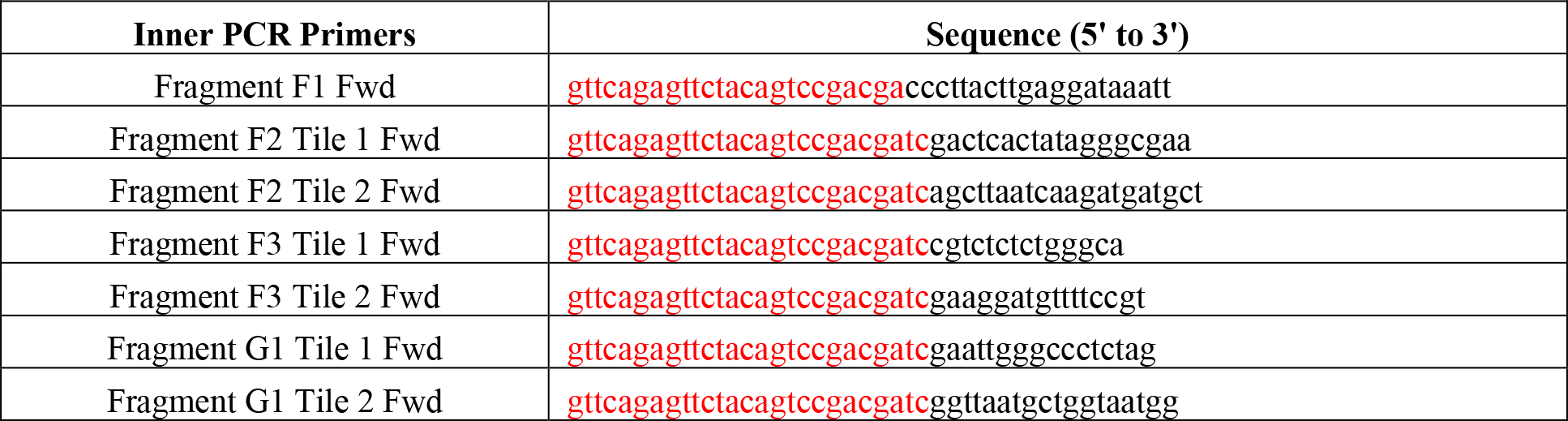
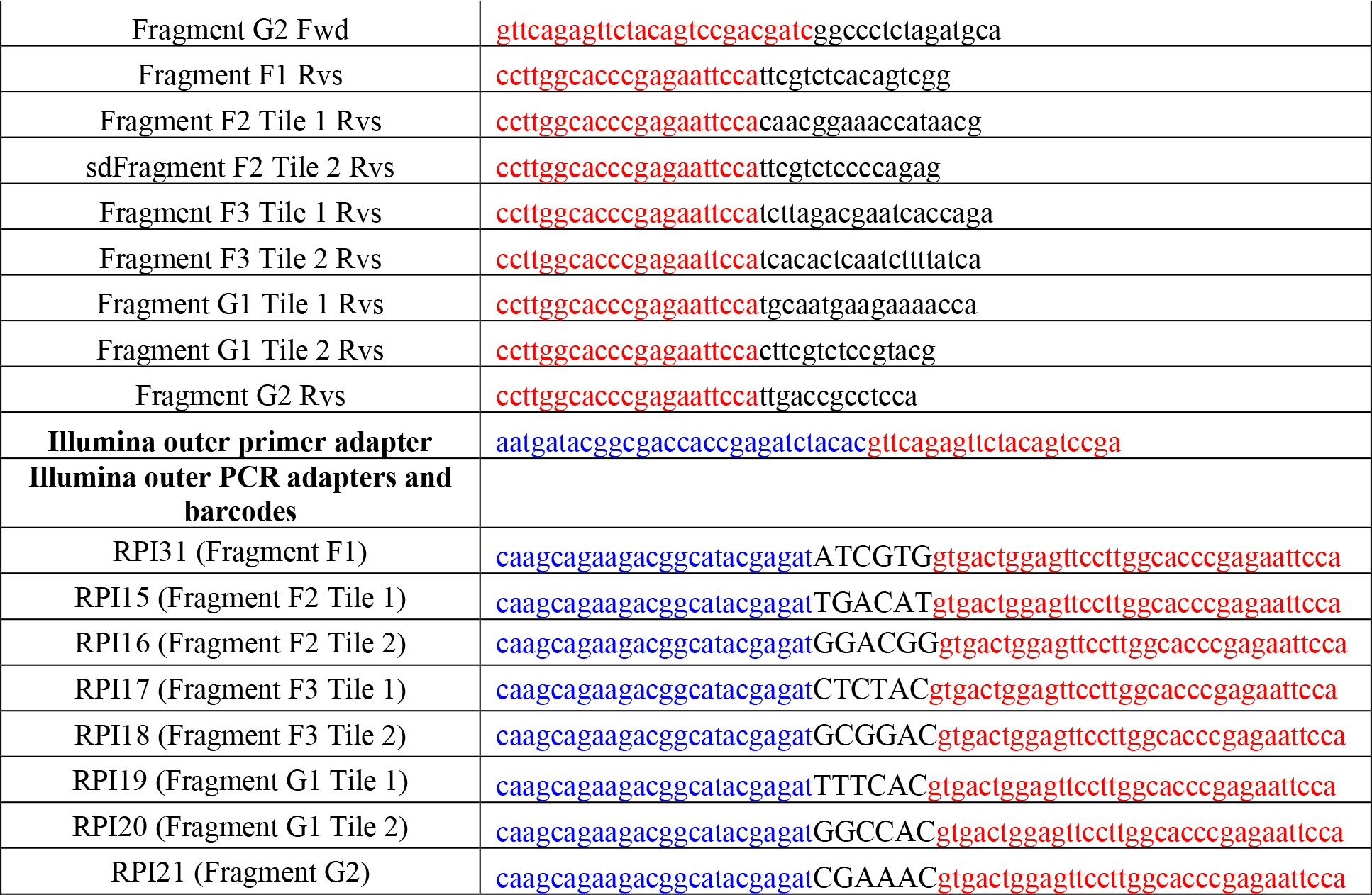
Inner and outer primers for PCR reactions for Illumina sequencing. Red indicates overhang regions for attaching Illumina adapter primers (inner PCR primers) or overhangs for attaching to inner PCR product (outer PCR primers), black is the overlap region in the gene or the barcode, blue is the Illumina adapter.

**Supplemental Table S3.**
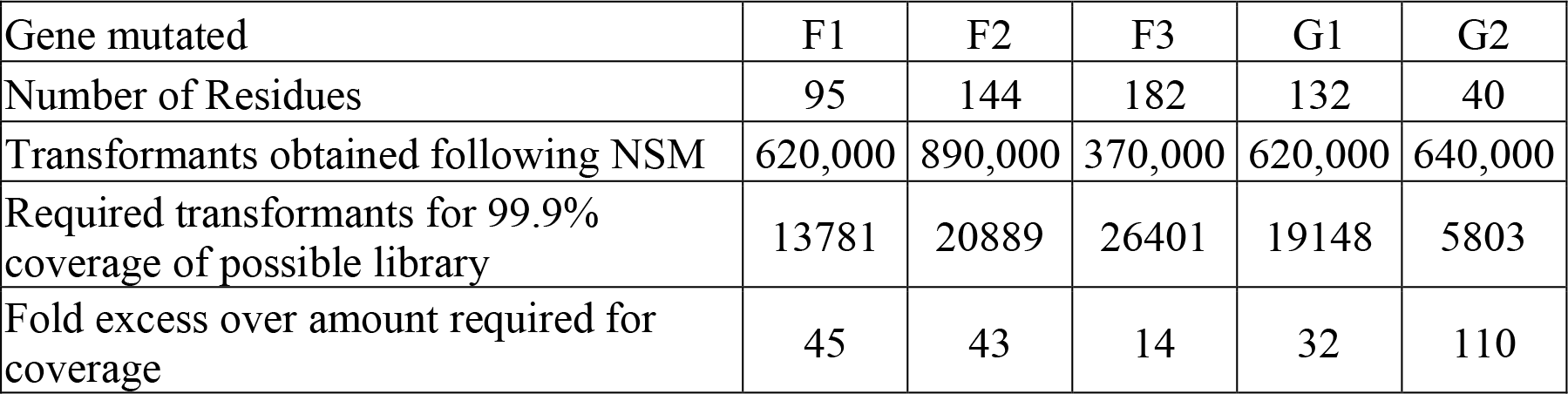
Mutant library preparation summary. Summary table of the transformants required for sufficient library coverage, transformants obtained during comprehensive mutant library preparation by nicking scanning mutagenesis, and the fold excess of the number of transformants required for coverage.

**Supplementary Table S4.**
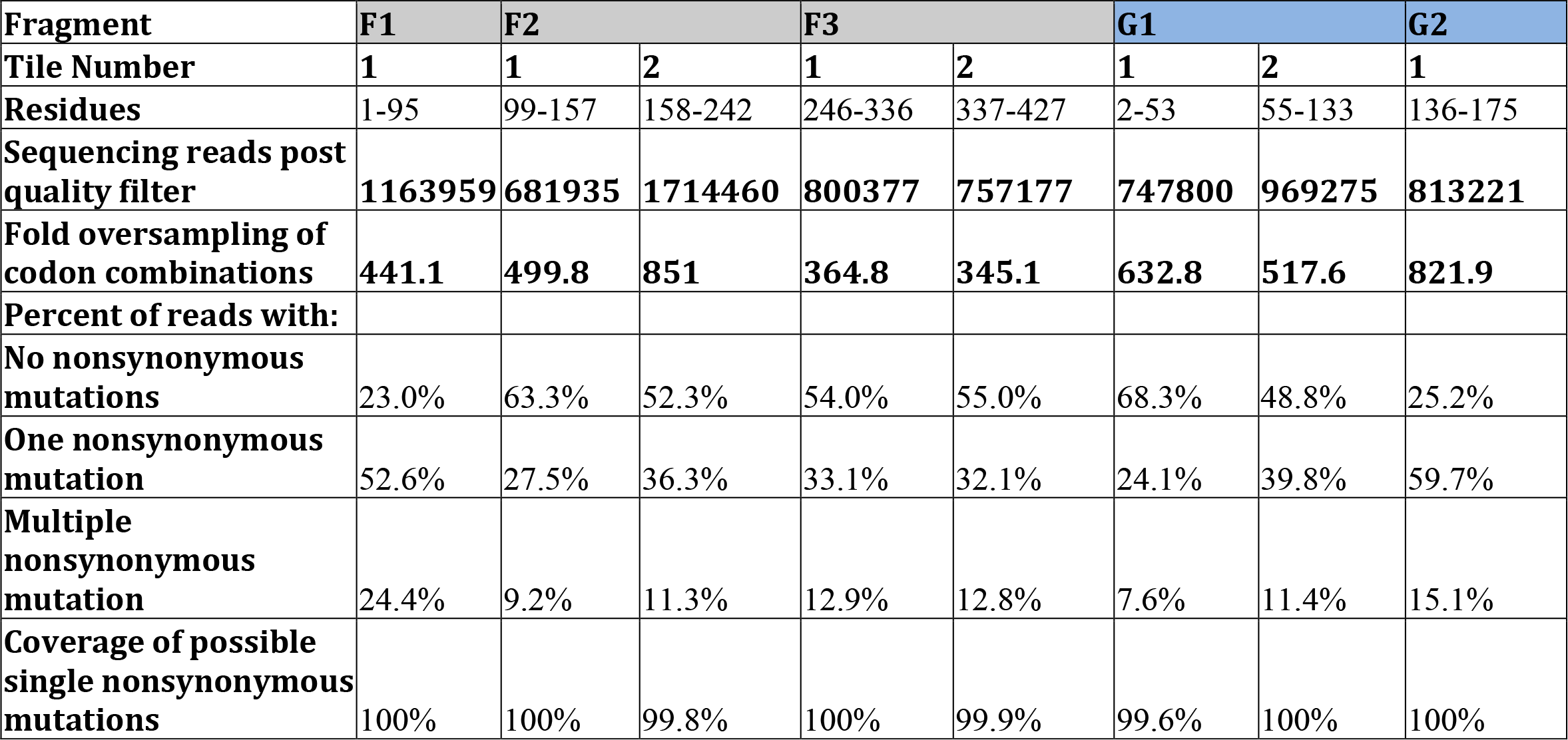
Mutant plasmid library NGS statistics. Summary table of the libraries prior to viral genome assembly. Long fragments required separate amplicon sequencing reactions as shown.

**Supplemental Table S5.**
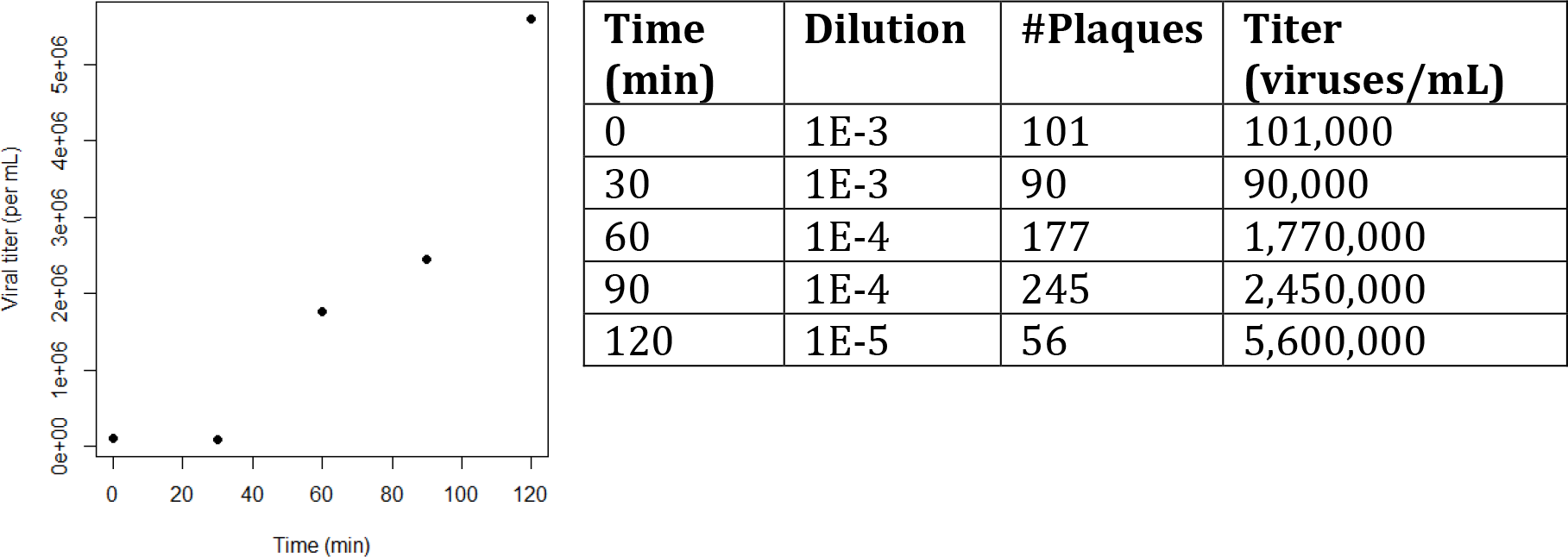
Viral growth rate during transformation cell recovery. Viral titers immediately after transformation (0 min) and then for every 30 min after up to 2 hours. As the XL1-Blue cells are not susceptible, the increase in titer is the result of cell bursts, which releases viable phage. While the virus is replicating for the first 30 minutes, they are all held within a cell and will only make one plaque.

**Supplemental Table S6.**
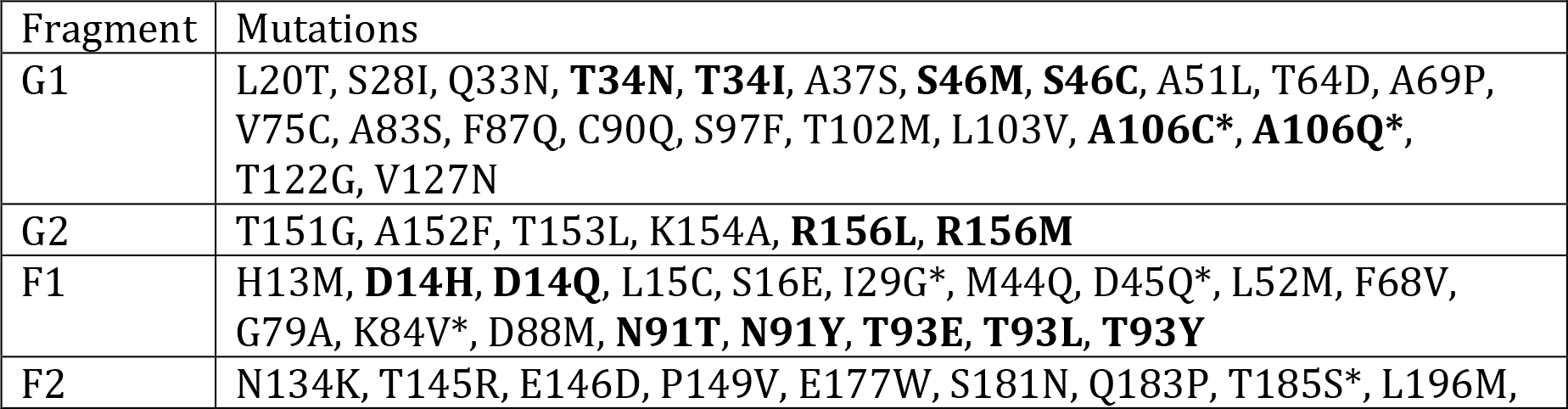
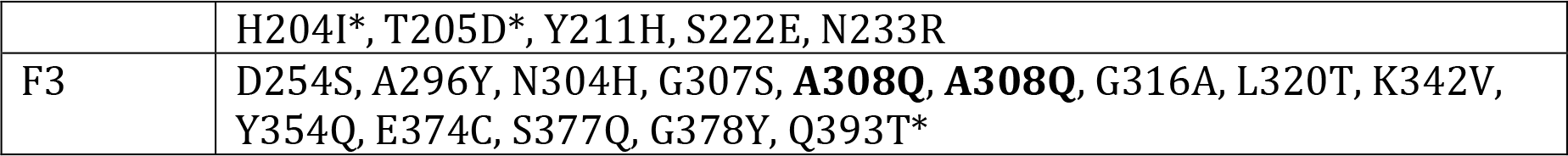
List of observed mutations in plaque sequencing. Residues where more than one substitution was observed are bolded. Residues where mutations have been observed in previous studies (but not the exact substitution) are indicated*.

**Supplemental Table S7.**
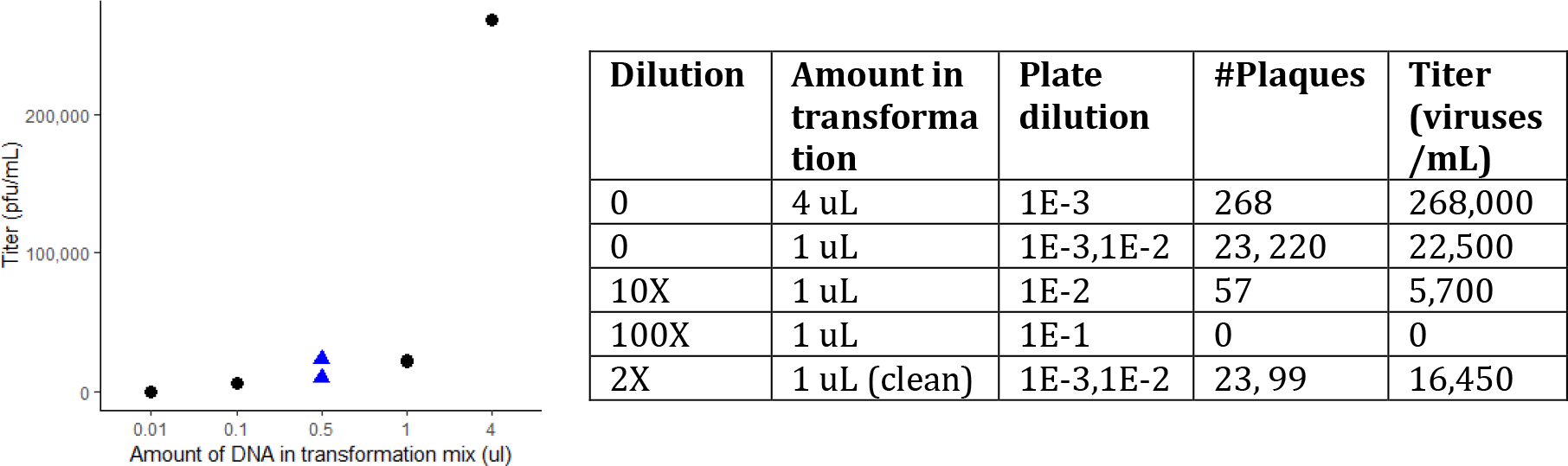
Testing for the optimal amount of ligation mix to transform. Different amounts and dilutions of the ligation mix were transformed into XL1-Blue cells to test for effects of DNA and salt concentration on transformation efficiency. The most transformants results from transforming 4 uL of ligated fragments.

